# Phase separation of hnRNP A1 upon specific RNA-binding observed by magnetic resonance

**DOI:** 10.1101/2022.03.21.485092

**Authors:** Irina Ritsch, Elisabeth Lehmann, Leonidas Emmanouilidis, Maxim Yulikov, Frédéric Allain, Gunnar Jeschke

## Abstract

Interaction of heterogeneous nuclear ribonucleoprotein A1 (hnRNP A1) with specific single-stranded RNA and its relation to liquid-liquid phase separation were investigated *in vitro* by magnetic resonance based on site-directed spin labelling. An ensemble model of free hnRNP A1 in the absence of RNA was derived from distance distributions between spin labelled sites and small angle X-ray scattering. This model revealed a compact state of the low-complexity domain and interaction of this domain with the RNA recognition motifs. Paramagnetic relaxation enhancement NMR spectroscopy confirmed this interaction. The addition of RNA to dispersed solutions of hnRNP A1 induced phase separation, observed by formation of liquid droplets. The phase separation depended on the RNA concentration and sequence, with continuous wave EPR spectroscopy showing that local protein dynamics is affected by point mutations in the RNA sequence. We propose that an interplay of sequence-specific RNA binding and phase transition serves as a regulatory mechanism for RNA segregation in the stress response of cells.

## Introduction

Liquid-liquid phase separation (LLPS) is a reversible de-mixing process that underlies the formation of membrane-less organelles in cells.^[1–5]^ These multi-component compartments play an important role in cell response to stress conditions^[6,7]^ and can be studied *in vitro* after isolation of the key components via the formation of liquid droplets (LDs). The main cellular mediators of LLPS are proteins that are equipped with multivalent, highly flexible protein domains, resp. inter-domain linkers,^[8]^ that are capable of driving LLPS *in vitro*.^[9-11]^ In this study, we focus on the human heterogeneous nuclear ribonucleoprotein A1 (hnRNP A1),^[6,7,12,13]^ which co-localizes with markers of stress granules under osmotic stress conditions.^[3,6–8]^ Among multiple functions of hnRNP A1 are alternative splicing regulation^[16]^ and mRNA export from the nucleus to the cytoplasm.^[17–20]^ In its central role for gene regulation, the function of hnRNP A1 has implications in many diseases, for example as a target for IRES-mediated viral infections,^[21–23]^ HIV viral infections,^[24–26]^ or cancer formation.^[27-31]^ Furthermore, it has been associated with neurodegenerative diseases, where point mutations in hnRNP A1 have been linked to a variant of inclusion body myopathy,^[32]^ as well as amyotrophic lateral sclerosis.^[32–34]^ HnRNP A1 consists of two homologous but functionally non-equivalent RNA-recognition motif (RRM) domains,^[35]^ which bind a large variety of RNA targets while maintaining some sequence specificity,^[12,36,37]^ and a carboxy-terminal glycine-rich domain (residues 197-320). The protein lacking the G-rich domain, but containing both RRMs (residues 1-196), is also known as unwinding protein 1 (UP1). The G-rich domain, due to its low compositional variation often referred to as ‘low complexity domain’ (LCD), does not adopt a fixed conformation in solution.^[9]^ In the context of structural modelling it can thus be classified as an intrinsically disordered domain (IDD). The LLPS properties of purified hnRNP A1 and other members of the hnRNP protein family were characterized in detail *in vitro*,^[9,11]^ where phase separation can lead to the formation of LDs, hydrogels, and insoluble aggregates.^[9,32,38]^ The G-rich domain of hnRNP A1 is necessary and sufficient to induce LLPS.^[9,39]^ and recently the importance of the regular spacing of aromatic residues was reported.^[39]^ In contrast, UP1 does not autonomously phase separate. However, structural characterization in the dispersed and phase separated state revealed the importance of largely, but not purely electrostatic interactions involving sites both in the G-rich domain and the RRMs during LLPS.^[40]^

Typically, phase separation *in vitro* is studied by making controlled perturbations of protein sample conditions, including changes in pH, ionic strength, temperature, or addition of small molecules like crowding agents.^[2]^ We here focus on studying interactions that drive LLPS in solutions of hnRNP A1 in the presence of RNA. Understanding the interactions in such mixed protein/oligonucleotide phase-separated solutions is receiving increased attention.^[10,41-44]^ For example it has been reported that RNA can modify LD morphology and viscosity,^[45–47]^ but little is known at the molecular level. We here study the interplay of RNA binding and phase separation of hnRNP A1, which is potentially relevant for regulation of splicing. To this end, we use specific short single-stranded RNAs derived from human intronic splicing silencer N1 from SMN2 gene transcript (SMN2-ISS-N1), a natural target sequence of hnRNP A1 involved in survival of motor neuron exon 7 splicing.^[12,48–51]^ RNA binding by UP1 has been studied in detail,^[36,51-56]^ and high-resolution structures of the individual RRMs as well as UP1 in complex with oligonucleotides are available.^[32,51,52,54,56]^ Here, we address RNA binding to full-length hnRNP A1, and the ensuing RNA-induced LLPS that we observe by complementary *in vitro* biophysical methods. In particular, we use site-directed spin labelling (SDSL) in combination with nuclear magnetic resonance (NMR) spectroscopy, pulsed and continuous wave (CW) electron paramagnetic resonance (EPR) spectroscopy. We localize the G-rich domain of full length hnRNP A1 in the dispersed state in the absence of RNA by paramagnetic relaxation enhancement (PRE) measurements,^[57,58]^ double electron-electron resonance (DEER) spectroscopy, and small-angle X-ray scattering (SAXS).^[59]^ The resulting solution state model of hnRNP A1 serves as a basis for interpreting RNA sequence dependence of LLPS upon addition of SMN2-ISS-N1-derived RNAs to dispersed hnRNP A1.

## Results and Discussion

For our *in vitro* study of RNA-induced LLPS of hnRNP A1 we recombinantly over-expressed and purified protein constructs from *E. coli*. cultures using a slightly modified version of the affinity-tag purification protocol with a cleavable His-tag reported previously (for details see SI).^[60]^ To understand the role of the C-terminal G-rich domain in RNA-binding, we compared full-length hnRNP A1 (most abundant isoform P09651-2, 320 residues) to ‘RRM-only’ UP1 (residues 1-196), see Figure 1(A,B). First, we characterized the structure of full-length hnRNP A1 in the absence of RNA in the dispersed state (<100 μM hnRNP A1 in 50 mM sodium phosphate, pH 6.5, 100 mM L-arginine, resp. L-glutamate) by applying a recently developed spin labelling approach for modelling of disordered protein domains^[61,62]^

**Figure 1.**
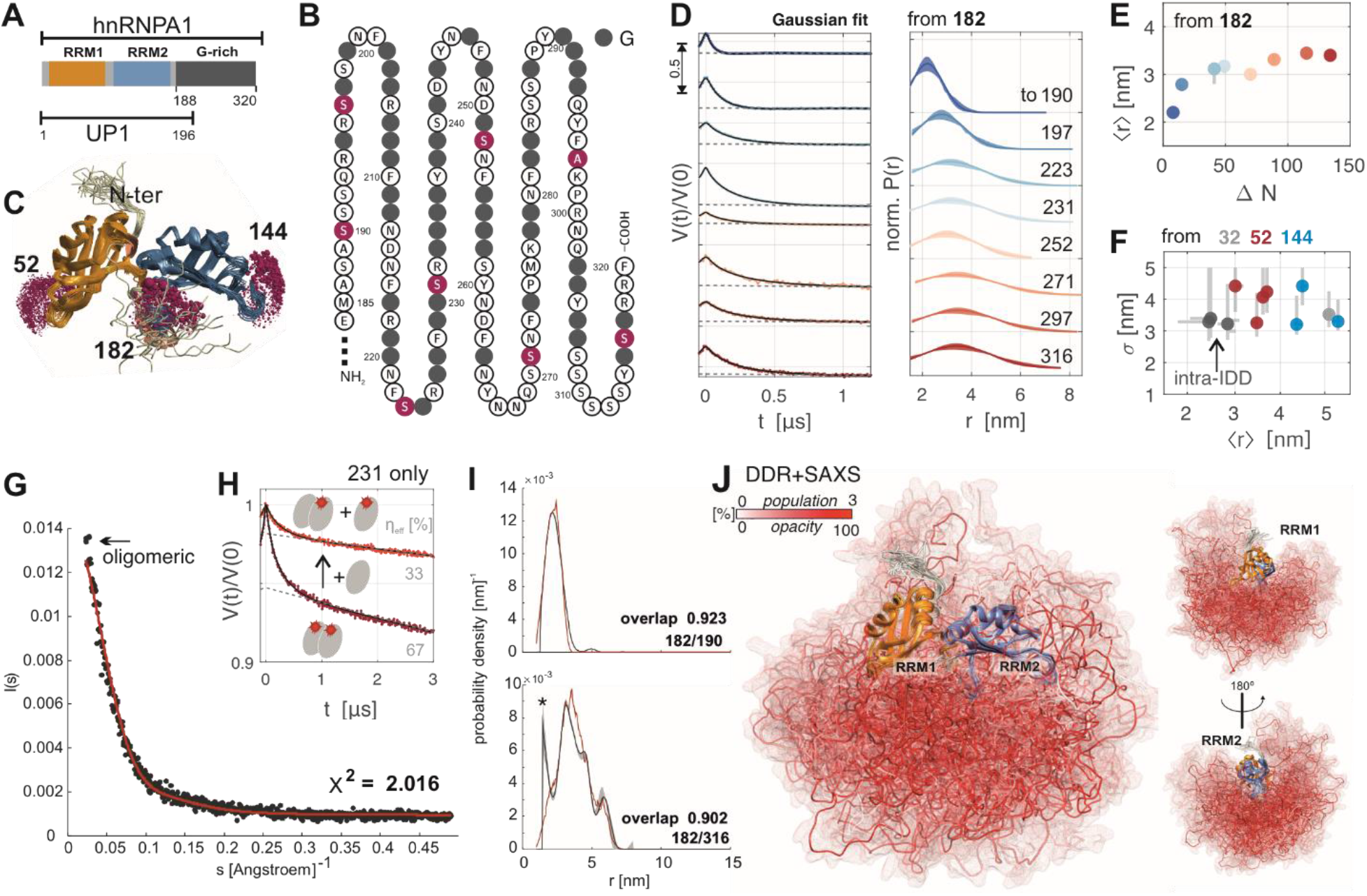
Ensemble model of hnRNP A1. (A) Domain organization of hnRNP A1 and UP1; (B) G-rich domain (197 to 320) of hnRNP A1, grey filled circles represent glycine, spin labelling sites are marked purple; (C) solution structure of UP1 [pdb: 2LYV] with spin-label rotamer simulation (purple spheres) at the major beacon sites; (D) primary data (left, arrow indicates modulation depth) and single Gaussian fits (right) of dipolar distance restraints measured from beacon 182 to sites in the G-rich domain; (E) mean distance 〈r〉 as a function of primary sequence separation ΔN of fits in (D); (F) width σ vs. 〈r〉 of remaining distance restraints, error bars from validation with DeerLab (https://jeschkelab.github.io/DeerLab); (G) ensemble SAXS curve and CRYSOL fit with DDR+SAXS model of hnRNP A1, the first data points were omitted in fitting to exclude contribution of oligomers; (H) primary DEER data (red), background fit (dashed grey), and fit (black) for hnRNP A1 spin labelled at site 231 before and after mixing with 1:1 unlabelled protein (total effective labelling efficiency η=c_spin_/c_protein_ indicated); (I) exemplary DDR fulfillment of the DDR+SAXS model; (J) conformer population-weighted visualization of hnRNP A1 structure ensemble model obtained with DDR+SAXS restraints.

We found that spin labelling of the two partially buried native Cys in the RRMs with commonly available nitroxide spin labels led to rapid, irreversible protein aggregation, most likely due to disrupting the native fold of the RRMs. Therefore, we designed variants with both native Cys mutated to non-reactive residues (C43S, C175A) as background in all SDSL experiments. Following established design criteria^[61]^ we engineered by site-directed mutagenesis solvent-accessible spin labelling sites that are remote from known RNA binding sites. These labelling sites in structurally well characterized UP1 serve as spatially well separated reference points (beacon sites) in the folded domains: 52 in the β2-β3 loop of RRM1, 144 in the equivalent β2-β3 loop of RRM2 (β7-β8 in full length hnRNP A1), and 182 in the C-terminal helix of RRM2 (see rotamer simulations with MTSL spin label in Figure 1 (C)). We also selected a total of eight ‘reporter’ sites for sampling the G-rich domain (190, 197, 223, 231, 252, 271, 297, and 316; all in this series except A297 are serines). This selection aimed at approximately uniform primary sequence intervals (see Figure 1 (B)) with minimal perturbation of inter-residue interactions by introducing cysteine. All mutants could be labeled well at approximately 25 μM hnRNP A1 and ten-fold molar excess of methanethiosulfonate spin label (MTSL, for details see SI). Doubly spin labelled hnRNP A1 mutants were used for distance distribution measurements by DEER spectroscopy (experimental details reported in SI). These measurements either involved one beacon site in an RRM and one reporter site in the G-rich domain (‘inter-domain restraints’), or two reporter sites within the G-rich domain (‘intra-domain restraints’). As controls, we also measured distance distributions for the pairs of beacon sites in UP1.

All 19 distance distributions involving at least one site in the G-rich domain were found to be very broad (see Figure 1 (D) for a selection), confirming the expected large flexibility of the G-rich domain. The shape of most distance distributions was approximated well by a two-parameter single-Gaussian distance distribution (mean distance 〈*r*〉 and distribution width *σ*).^[62]^ The detailed data analysis protocol, including validation of fitting, can be found in the SI. Our previous methodological study revealed that deviations from a Gaussian distribution are insignificant compared to other uncertainties in ensemble modelling.^[62]^ Hence, we can discuss the general trends in terms of 〈*r*〉 and σ. We first tested whether the G-rich domain can be considered as a random coil. This would imply that 〈*r*〉 and σ increase monotonously with increasing primary distance separation Δ*N*. In particular, 〈*r*〉 should scale as Δ*N^V^* with *v* = 3/5 in the good solvent case and *v* = ½ if interactions of the chain with the solvent are as strong as within the chain (θ solvent). For the beacon site 182 close to the N terminus of the G-rich domain, Δ*N* directly relates to an effective end-to-end segment length in this domain. As seen in Figure 1(D,E,F), the increase of 〈*r*〉 and σ conforms to this expectation only for Δ*N* < 50. The distance with Δ*N* 70 (182/252) is even slightly shorter than distance with Δ*N*=49 (182/231), although this difference is small compared to the large distribution widths. In any case, for the remaining distance distributions with Δ*N*>89 mean distance and width do not increase significantly.

For the distance measurements from other beacon sites 52, resp. 144 to reporter sites in the G-rich domain (red, resp. blue in Figure 1 (F)), as well as for distance measurements between two reporter sites in the G-rich domain (intra-IDD, grey in Figure 1 (F)), we consider the relation of the distribution widths σ to the mean distance 〈*r*〉 (for primary data and fits see SI). We find that the reporter sites in the G-rich domain are on average closer to the loop of RRM1 (represented by beacon 52, red) than to the equivalent loop in RRM2 (beacon 144, blue). The G-rich domain thus has a slight preference for an asymmetric placement with respect to the RRMs, yet this trend is weak, given that all distance distributions are very broad. Second, despite a variation of Δ*N* between 40 and 85 the intra-domain segments (grey) in the G-rich domain (231/271, 231/316, 271/316) all have similar short mean distances, and are surprisingly narrow. This indicates that the G-rich domain mostly samples rather compact conformations.

For a more detailed analysis, we used the distance distribution restraints (DDRs) for constructing an atomistic structural ensemble model of full length hnRNP A1 using a previously reported force-field free algorithm.^[61–63]^ The individual conformers in the ensemble model are constructed by *in-silico* growing the domain from a chosen anchor residue at the C-terminal end of the solution model of UP1 (pdb 2LYV).^[60]^ As the C-terminal section of UP1 (188-196) is only weakly experimentally restrained and strongly diverging,^[60]^ we constructed conformers for residues 188-320. A large stochastic raw ensemble of individual conformers of the G-rich domain was generated using the 19 DDRs as Gaussian restraints, and taking into account the primary sequence by sampling from residue-specific Ramachandran-statistics.^[61]^ This ensemble was refined and contracted using the EnsembleFit module of MMM, additionally taking into account an experimental SAXS curve (Figure 1 (G)) (experimental details see SI).^[63]^ This step fits conformer populations and discards lowly populated conformers, while aiming for a balanced fit to the DDRs and SAXS curve. For SAXS fitting with CRYSOL,^[64]^ the very short angle (*s* < 0.02195 Å^-1^) region was truncated, since it is highly sensitive to distortions from weak contributions of aggregates or transient oligomers (marked ‘oligomeric’ in Figure 1 (G)) in the sample. The presence of such oligomers was confirmed by DEER measurements with diamagnetic dilution (Figure 1 (G)). For population fitting, we specified DDRs by model-free distributions obtained by Tikhonov regularization. Including jack-knife resampling and a second round of population re-weighting,^[62]^ we arrived at the final structural ensemble of full length hnRNP A1 consisting of 138 conformers visualized in Figure 1 (J). The opacity of the conformers in the visualization reflects their population in the refined ensemble. Figure 1 (I) shows by two examples that the input DDRs are well reproduced by the refined ensemble (see SI for complete data).

The ensemble model conforms that the G-rich domain is compact yet features a broad range of molecular conformations. It occupies space close to the linker region between the RRMs, distant from the N-terminal residues and close to the RNA-binding interface. The unexpectedly short distances measured between the RRMs and the C-terminal region (close to reporter site 316) can only be fulfilled by the G-rich domain on average looping back towards the RRMs. We checked that an ensemble based on only Ramachandran statistics and the SAXS curve does not significantly sample such compact conformations (see SI Figures S14 and S15). This may explain why a previous study by SAXS and coarse grained molecular dynamics-based simulations approach^[40]^ did not reveal the localization of the G-rich domain that we find here. The broad distribution of conformers in our model suggests an only slightly rugged energy hypersurface. This is in agreement with the transient interaction between the RRMs and the flexible G-rich domain found in the previous study.^[40]^

As mentioned above, the SAXS curve indicates presence of oligomers, but only to a minor extent. Since such oligomers might also affect RNA-binding experiments, we used the DEER experiment with singly-spin labelled hnRNP A1 to independently confirm and monitor their presence, by analyzing the ‘background’ inter-spin distance contribution.^[59,65,66]^ This contribution is sensitive to local spin label and thus local protein concentration. In Figure 1(H) we show DEER data obtained with for hnRNP A1 singly spin-labelled at site 231. The bulk of the data is fitted well by a slow decay (dashed grey line), which accounts for the expected dispersed, homogeneously distributed hnRNP A1 at the low micro-molar concentration used in our DEER experiments. Additionally, we observed a fast-decaying component with a small fraction of the modulation depth expected for a doubly spin-labelled entity. This confirms that a small fraction of protein is localized in a more concentrated, or ‘dense’ environment. Interestingly, when admixing unlabeled protein, i.e. when reducing the effective labelling efficiency *η*_eff_ = *c*_spin_/*c*_protein_, we observed substantial suppression of the steep contribution, thus a smaller contribution of the dense phase. This reveals that exchange between protein fractions in the dense phase and the admixed unlabelled protein must have occurred. We conclude that the interactions leading to the dense phase contribution must be at least partially reversible. If we assume that hnRNP A1 is distributed homogeneously at higher concentration in this dense phase, the observed modulation depth indicates that less than 5% of hnRNP A1 molecules are located in this phase.

In addition to identifying a small amount of dense phase in solutions of predominantly dispersed hnRNP A1, the singly spin labelled hnRNP A1 samples also allowed to validate our structural model by PRE experiments. In Figure 2(A) we show normalized PRE intensity ratios obtained from hnRNP A1 that is both spin-labelled and ^15^N-isotope-labelled (NMR active/EPR active). MTSL was placed at one of the three sites 231, 271, or 316. In these all-interaction experiments, we cannot distinguish *a priori* between PRE arising from inter- and intramolecular proximity of the label to a given residue. However, the DEER-based analysis indicates an only small fraction of molecules in the dense phase. We further performed intermolecular-only control experiments on a mixture of hnRNP A1 with the spin label in natural isotope molecules and ^15^N-isotope-labelled molecules without spin label (see SI). These experiments confirm that the intermolecular contribution is weak under the conditions that we use. By mapping the PRE patterns to the solution-state model of UP1 (Figure 2 (B), for experimental and analysis details see SI), we found moderate to strong signal reduction for a large fraction of the RRM surface. This is in agreement with the G-rich domain being disordered, allowing the flexible chain to sample many remote surface interaction sites. With the spin label at site 231, we observed clusters of residues in the N- and C-terminal flanking regions of RRM1, as well as in the β2-β3 loop and C-terminal helix of RRM2. Both regions constitute major sites for RNA-recognition. Most strikingly, significant backbone PRE effects were observed for R88 and V90, (RRM1), resp. R179 and R181 (RRM2), which have been reported by Beusch *et al*.^[51]^ to directly recognize the ‘AG’ motifs in SMN2-ISS-N1-related RNA variants by hydrogen bonding via the protein backbone. Thus, in the dispersed, RNA-free state, sites that form the RNA-binding interface coincide with sites that interact transiently with the G-rich domain around site 231. A similar PRE pattern was observed with the spin label at site 271, whereas a non-specific pattern was observed with the label at site 316. Using a recently published PRE-prediction algorithm,^[67]^ we see that the PRE pattern predicted from our structural ensemble model is in reasonable agreement with the experimental PREs (r.m.s.d.PRE = 0.0822, correlation shown in Figure 2 (C)), despite the PRE having been measured at ambient temperature, while the DEER data were measured with flash-frozen samples.

**Figure 2.**
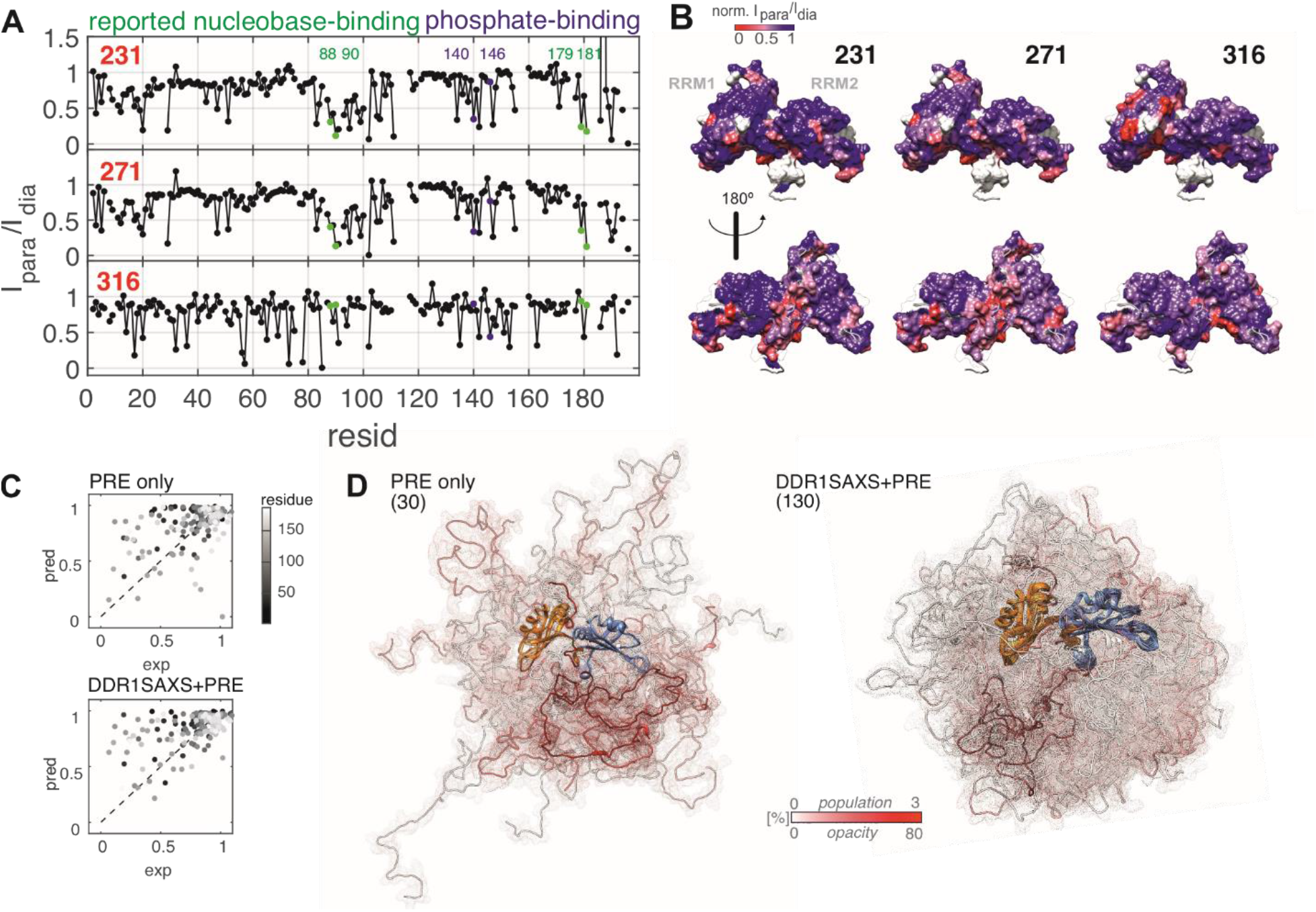
Validation of the ensemble model and determination of RRM sites that interact with the G-rich domain by PRE experiments. (A) PRE experiments reported as relative cross-peak volume ratio I_para_/I_dia_ in ^1^H-^15^N-HSQC obtained with 50 μM hnRNP A1 spin labelled at sites 231, resp. 271, resp. 316, data were normalized to peak-volume-ratio at residue 117 (buried). Labelled green, resp. blue dots indicate backbone nitrogens that directly coordinate RNA bases, resp. RNA backbone as identified in Beusch et al.;^[51]^ (B) mapping of PRE to the surface of UP1; (C) PRE predictions vs experimental values based on a previously unrefined model (‘PRE only’), resp. on the DDR+SAXS+PRE refined ensemble model; (D) ensemble models used for analysis in (C); RRM1 is shown on the left, and the number of chains that remain after population reweighting is indicated. Transparency and color of G-rich domain chain reflects conformer population.

Going one step further, we included the PRE data at the ensemble refinement stage of our modelling. Thereby, PRE fit quality could be improved (r.m.s.d._PRE_= 0.0446) while simultaneously maintaining high fit quality for DEER and SAXS data (Figure 2(C), and SI Table S4). We attribute the still mediocre PRE fit quality to the influence of backbone dynamics on PRE that we neglected. Interestingly, an ensemble fitted to only PRE restraints from a raw ensemble considering only residue-specific Ramachandran statistics resulted in a slightly worse PRE fit (r.m.s.d._PRE_ = 0.0581). Comparison of this PRE-only ensemble with the DDR+SAXS+PRE ensemble (Figure 2 (D)) reveals that, using the same number of conformers in the raw ensemble, DDR-based conformational sampling found a significantly larger number of conformers that are consistent with the PRE data. These conformers are simultaneously consistent with the DEER and SAXS datasets.

We were intrigued by the overlap between RNA-binding sites and the residues for which MTSL at sites 231 or 271 induces strong PRE. Shielding of the RNA binding face would imply that the G-rich domain is displaced upon RNA binding. This is in line with a model of bound RNA proposed earlier by Beusch et al.,^[51]^ which assumes an RNA with the wild-type SMN2-ISS-N1 sequence (Figure 3A). Such RNA can bind to UP1 in a 1:1 ratio, with RRM1 binding a 3’-AG motif, and RRM2 binding a 5’-AG motif. In this complex, the RNA wraps around both RRMs and the linker region, covering a large fraction of the positively charged surface of UP1 (visualized in Figure 3 (B) with Coulombic color-coding of the solvent accessible surface). With our ensemble model of free hnRNP A1 in hand, we investigated such a potential role of the G-rich domain in RNA binding. We used four RNA variants where either the 5’ motif (‘RNA*ag*’) or the 3’ motif (‘RNA*ga*’), or both (‘RNA*aa*’) have been mutated in the same manner (in this notation ‘wild type’ corresponds to ‘RNA*gg*’). This choice is motivated by the earlier finding that already a single point mutation of these two core 5’-AG-’3 binding motifs (AG→AA) in the RNA sequence can strongly affect the splicing repression by hnRNP A1.^[51]^ In such purine-to-purine point mutations, adenine as the substituting nucleobase has similar overall size and aromatic properties as the wild-type guanine, yet guanine has a higher hydrogen bonding capacity.

**Figure 3.**
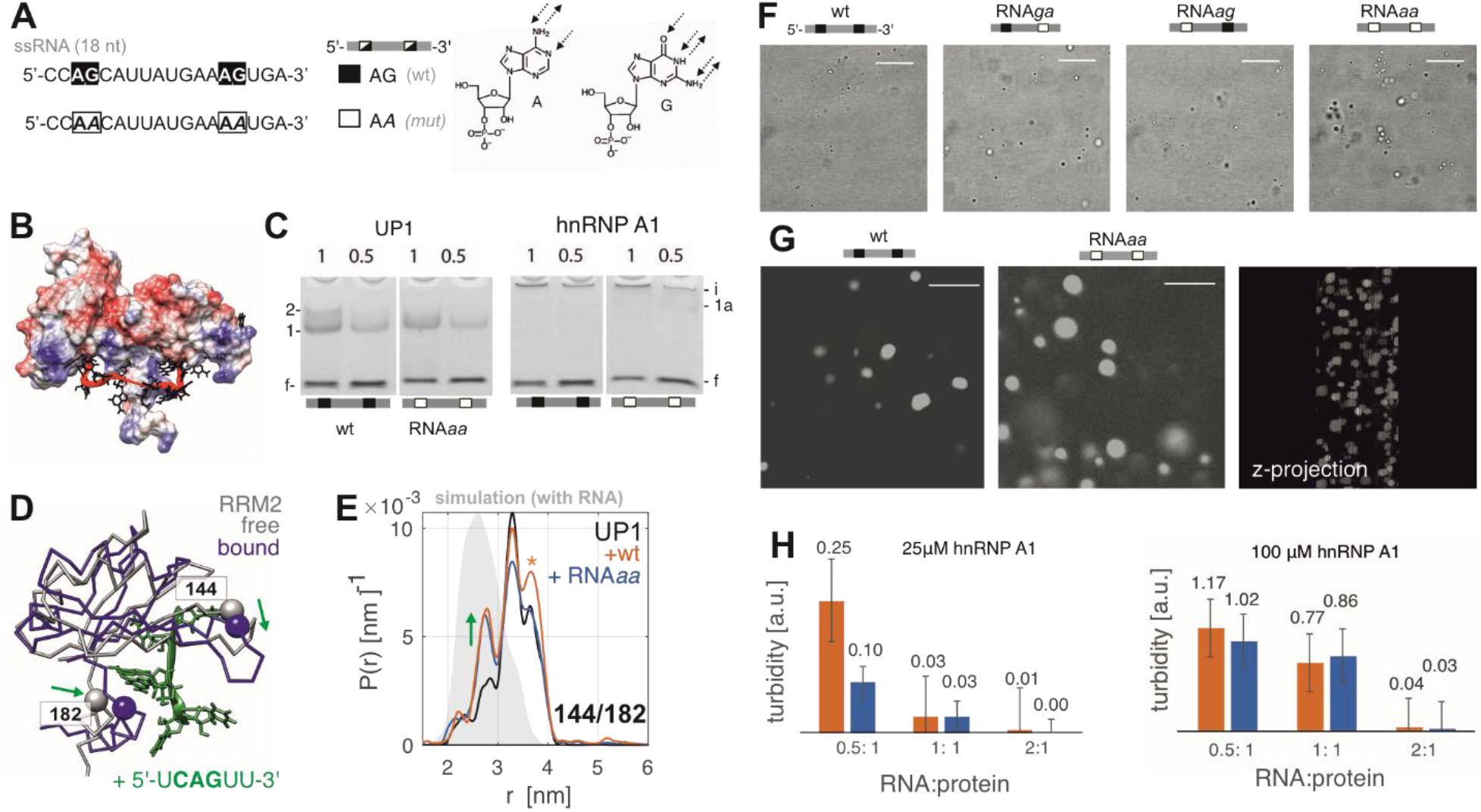
Effects of RNA binding to hnRNP A1. (A) ssRNA variants derived from SMN2-ISS-N1 wt sequence with one, resp. two G→A point mutations; (B) Coulombic colored model of RNA-bound UP1 from Beusch et al.;^[51]^ (C) toluidine-stained EMSA of 50 μM wt resp. doubly mutated RNAaa incubated with UP1 resp. hnRNP A1 at two stoichiometric ratios of protein relative to RNA as indicated above the lanes; f free RNA, i non-migrating material in the gel pocket, other bands are enumerated; (D) visualization of the change in UP1 conformation upon RNA binding expected from the solution structure of UP1 [pdb: 2LYV] and the model of RNA-bound UP1 from Beusch et al.;^[51]^ (E) DEER distance distribution between beacon sites 144/182 of free UP1 (black), UP1 + 1:1 wt RNA (orange), and UP1 + 1:1 RNAaa (blue) overlayed on the MMM simulation with free UP1 (grey area), resp. RNA bound RRM2 (green area, pdb: 5MPL); (F) confocal transmission microscopy images of 25 μM hnRNP A1with 1:1 RNA variants, scale bar in all panels of this figure is 10 μm; (G) left, center: fluorescence microscopy images of 25 μM hnRNP A1 with 0.5:1 RNA:protein from two RNA variants stained with GelRed^®^ RNA stain. right: projection of fluorescent image stack in third dimension obtained with 1:1 RNA:protein ratio and RNAaa (H) turbidity determined by OD_600_ of 100 μM, resp. 25 μM hnRNP A1 with wt RNA (orange) and RNAaa (blue) at indicated ratios.

RNA binding to UP1 in our buffer conditions was confirmed by non-denaturing electrophoretic mobility shift assays (EMSA, see Figure 3 (C)). With RNA-staining by toluidine, we observed a resolved band with reduced mobility compared to free RNA upon 1:1 stochiometric incubation of UP1 with either wild-type RNA. The same band was observed with RNA*αα*, showing that even the doubly mutated RNA can still bind to the RRMs. Because significant amounts of free RNA were observed, both at stochiometric ratio and at RNA excess, we concluded that the overall binding affinity is rather low with both wt and mutated RNA. A faint second resolved band was observed with wt RNA, which is likely related to a RNA/protein complex of different stoichiometry. The absence of this second band with mutated RNA might be related to different dynamics of binding and unbinding of the mutated RNAs, which could lead to broadening of the second band to an extent that makes it invisible. In contrast to the mobile bands observed with UP1, EMSA with the same RNAs and hnRNP A1 led to the accumulation of immobile material in the gel pockets (annotated (i) in Figure 3 (C)). A very faint band of a slowly migrating complex is visible with RNA*aa*, but not with wt RNA. As is the case with UP1, significant amounts of free RNA are also observed with hnRNP A1. Thus, while some RNA is trapped in the aggregates with hnRNP A1, a substantial amount remains unbound.

In order to identify the immobile component in EMSA, we used confocal microscopy and optical turbidity measurements at 600 nm, which are well established methods to monitor aggregation and LLPS.^[9,68]^ The samples were stabilized by a low melting agarose gel (1% w/v), which prevents LD sedimentation and allowed stable observation of the samples for several hours.^[68]^ We observed LLPS upon adding RNA to 25 μM hnRNP A1 (shown at 1:1 ratio) by confocal microscopy as the formation of many small LDs (~1 μm) as shown by confocal microscopy in Figure 3 (F). In addition to LDs, recognized by their spherical shape and occasional fusion events, we sometimes observed non-spherical particles and insoluble aggregates (see SI). All four RNA variants were able to induce LLPS with hnRNP A1 and none of them with UP1 (see SI). The size and appearance of the agarose-stabilized LDs depended on absolute protein concentration and RNA-to-protein ratio. Large LDs (up to 10 μm) are formed upon addition of wt RNA, resp. the doubly mutated RNA*aa* at a 0.5:1 (RNA:protein) molar ratio to 100 μM hnRNP A1, as seen by fluorescence microscopy when incubating with GelRed® RNA-stain (Biotium, USA) fluorescent dye (Figure 3(G)). The dye appears to be distributed homogenously within each droplet, as can be seen in side-view images from reconstructing focal plane stack, suggesting a homogeneous distribution of RNA in the droplets. The turbidities for the conditions used for imaging as well as for additional RNA:protein ratios are shown in Figure 3(H). Surprisingly, at an hnRNP A1 concentration of 25 μM, we observed an RNA-sequence dependent LLPS effect, where RNA-induced LLPS was weaker for the mutated RNA variants than for the wt RNA (Figure 3(H)). To understand how such RNA-sequence specific effects could be mediated in full length hnRNP A1, we screened for conformational changes upon binding by different RNA variants to UP1, where we know that no LLPS occurs. Based on structural information of RNA bound to the folded domains,^[51,56]^ we selected the pair of beacon sites 144 and 182 in RRM2 for this test. Comparison of the conformations of RRM2 in the absence of RNA and in complex with a short RNA containing the core motif of wt RNA^[51]^ (Figure 3(D)) suggests that, upon binding the C-terminal helix moves towards the RRM2 β2-β3 loop region. Indeed, we observe a shift of conformer distribution that is even more pronounced than expected from simulations with the NMR structures of free UP1 and of the RNA-RRM2 complex Figure 3(E)). However, the free state was already found shifted to longer distances than expected from modelling, which somewhat limits our interpretation. We did not find substantial differences in the bound contribution between the wt and doubly mutated RNA.

In both cases, only a fraction of the distribution shifts to shorter distances. In the case of wt RNA, we observed a slightly enhanced dipolar modulation amplitude which we attribute to RNA-enhanced protein-protein contacts, in agreement with the weak second band observed in EMSA. The upper distance shoulder of the distribution is slightly increased with wt RNA (marked by asterisk in Figure 3(E)), which is likely to be an artefact of the enhanced spin interactions due to multi-spin effects in those additional complexes.^[69]^ Overall, the DEER binding analysis to the RRMs did not provide a clear explanation of RNA-sequence dependent LLPS at low protein concentration. Unfortunately, the phase-separated state that is induced by RNA binding to full length hnRNP A1 has a too high spin concentration for analogous distance distribution analysis (see SI Figure S7).

Therefore, we turned to ambient-temperature CW EPR spectroscopy to investigate dynamic changes of the G-rich domain upon RNA binding and LLPS. We expected that local molecular crowding would lead to reduced backbone and spin label side chain mobility. Such reduced mobility is observable by line broadening in CW EPR spectra of singly spin-labelled mutants, which reports on dynamics in the high pico-to low nanosecond range. The results of this dynamics study for reporter sites 231, 271 and 316 in the G-rich domain are presented in Figure 4. Additional results for labelling sites in the RRMs and for UP1 can be found in the SI. We first characterized the dynamics in the RNA-free, dispersed reference state. The three spectra in agarose-containing buffer, analogous to imaging conditions in Figure 3, show the expected very fast spin label mobility (black lines in Figure 4(E)). The lineshapes could be fitted well using the Easyspin^[70,71]^ toolbox, assuming a single isotropically tumbling component with rotational correlation time τ_corr_ (fits and fitting procedure shown in SI). Site 316, close to the C-terminus, has the shortest correlation time and thus highest mobility (τ_corr_ = 0.35 ns), followed by site 271 (τ_corr_ = 0.53 ns) and site 231 (τ_corr_ = 0.60 ns). We expect that sidechain mobility is governed by an interplay of local density and compositional bias that affects backbone dynamics. To understand the observed slight local mobility differences, we investigated whether the spin label mobility correlates with local sequence context and average ensemble density at the label site.

**Figure 4.**
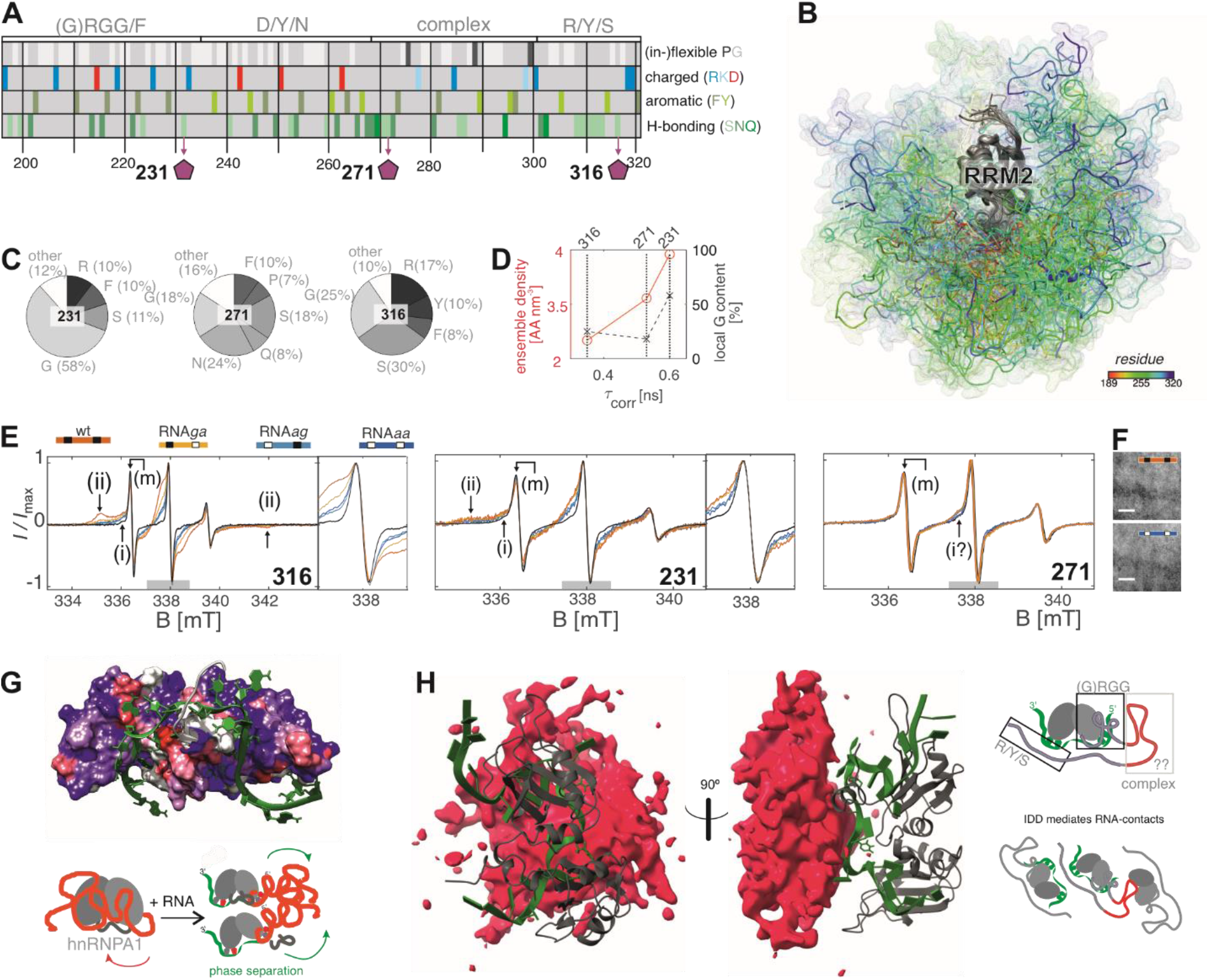
Relation between RNA binding and LLPS. (A) sequence breakdown of G-rich domain of hnRNP A1; (B) side view looking at RRM2 of full length hnRNP A1 ensemble model ‘DDR+SAXS+PRE’; G-rich domain chains are color-coded by residue (C) local amino acid type statistics in radius of 1 nm around reporter sites in model shown in (B); (D) fitted τ_corr_ of nitroxide spin label at reporter sites 231, 271 and 316 spectra vs. local amino acid composition, resp. vs. local ensemble density; (E) overview and zoom on central line of CW EPR of 25 μM hnRNP A1 reporter sites overlayed at 9.5 GHz, normalized to maximum intensity; black: free protein (no RNA); color-coded:+ 1:1 molar ratio of RNA variants (F) confocal image of hnRNP A1 mutant labelled at site 271 with 1:1 RNA, scale bar 10 μm; (G) RNA mapped on surface of RRMs color-coded by PRE from site 231 (stronger PRE shown as red, see Figure 2(B)) and scheme of proposed RNA vs G-rich domain competition mechanism for interaction with the RRMs; (H) RNA mapped on RRMs in ensemble model of hnRNP A1 with G-rich domain (red) displayed as average occupied volume, schematic drawing of the proposed cooperation mechanism (recognition of RRM-bound RNA).

Conceptually, such analysis in terms of local amino acid composition is related to the proximity to ‘stickers’ (interacting) and ‘spacers’ (stretches of non-interacting)’ amino acids in the primary sequence.^[11]^ Selected amino acid categories are schematically represented in Figure 4(A). Glycine and proline are highlighted in the top row and are expected to be key regulators of local high resp. low backbone flexibility. Below, we indicate different types of ‘stickers’: in the second row charged residues; in the third row aromatic residues; and in the last row hydrogen-bond donor and acceptor amino acids. With our structural ensemble model of full length hnRNP A1 at hand, we extended the local compositional analysis beyond primary sequence statistics, by including spatial aspects. To this end we calculated the average amino acid density and composition in a one nanometer spherical zone around the reporter sites from the structural ensemble model of free hnRNP A1 shown in Figure 4(B) (DDR+SAXS+PRE ensemble model, for computational details, see SI). The resulting local amino acid composition statistics are shown in Figure 4 (C) for amino acid types more abundant than 5%. The abundance of glycine, as well as the local average ensemble density are plotted against the reporter site mobility in Figure 4 (D). Site 231 with the highest τ_corr_ and thus lowest sidechain mobility is located in the densest region of the disordered domain (3.78 AA nm^-3^), but somewhat unexpectedly has the highest excess of flexible over inflexible residues (58% G, 0% P). In full agreement, with expectations, site 316 with the lowest density (3.03 AA nm^-3^, 25 % G, 0% P) has the highest experimental mobility. Site 271 falls in the intermediate mobility and density range, and is located at a site of higher sequence complexity, in particular close to proline (3.29 AA nm^-3^, 18% G, 7% P). On this small dataset the average ensemble density thus appears to be suitable for predicting local dynamics, and is somewhat more robust than individual side chain statistics. More detailed understanding might be achieved with additional dynamic ensemble modelling, but note that the static analysis presented here provides substantial insight at low computational effort for ensemble generation and analysis.

Having established spin label dynamics at the reporter sites for the free reference state, we studied changes upon mixing 25 μM spin-labelled hnRNP A1 with wt RNA, resp. doubly mutated RNA*aa*, which we know from turbidity measurements and microscopy to induce LLPS. We observed strongly broadened spectral components at site 231, indicating strong spin label mobility changes. Co-existence of multiple resolved spectral components implies that different populations of protein (e.g. dispersed vs RNA-bound and/or phase separated) co-exist in the sample, which interchange on a time-scale slower than the EPR experimental resolution of about 1 μs. In our case, dispersed protein with high sidechain mobility appears to coexist with phase-separated protein with lower mobility. The ratio of the mobile to the immobile component initially increases with the ratio of added RNA:protein, but plateaus after a ratio 1:1 (see Figure S11). We annotate rapid rotational tumbling, as also observed in the free state, as the mobile (m) component, moderate spectral broadening as the intermediate (i) component, and the contribution where the low-field minimum of the immobile nitroxide spectrum can clearly be identified as the immobile (ii) component.

Broadened spectral components were observed with all four RNA variants for the label at sites 231 and 316, but not at site 271. The latter appears to be a special case, since placing the spin label at this site substantially suppressed RNA-induced LLPS, as could be observed by the absence of LDs in imaging (Figure 4(F)), and by reduced turbidity compared to wild-type hnRNP A1 (0.036 ± 0.059 with wt RNA, and 0.004 ± 0.004 with RNA*aa*, at 0.5:1 RNA: protein ratio, for wild-type values see Figure 3(H)). This suppression is unexpected, since early work^[9]^ showed that LLPS of RNA-free hnRNP A1 did not require the presence of a ‘steric-zipper’ motif, located just N-terminal of site 271. In contrast, our study suggests that, in the context of RNA-induced LLPS, this region of the G-rich domain is critical for promoting phase separation. This is interesting, since 271 is located in a region that has a low G content and does not contain RGG motifs known to be involved in RNA binding, yet is enriched in amino acids with high H-bonding capacity (50 %= N/S/Q). As the RRM domains in all three constructs are the same, the observed perturbation must arise from positioning the spin label at different sites in the G-rich domain.

The CW-EPR spectra with labels at sites 231 and 316 agree with the previously discussed RNA-sequence dependent LLPS effect at low hnRNP A1 concentration that we observed by turbidity measurements. At site 231, the 5’ AG motif (present in RNA*ga*, and wt, but not in RNA*ag* and RNA*aa*) was found to be necessary and sufficient to observe an immobile component (ii). This implies that LLPS in our set of RNA variants is governed not only by the number of G nucleotides or AG di-nucleotide motifs; it appears to be sequence-specific to some extent. With RNA*ag* and RNA*aa*, a immobile component (ii) was not observed and the contribution of the mobile component (m) remained higher. That RNA-binding affects mobility at site 231 might be expected even in the absence of LLPS, since it is located close to an arginine residue and just C-terminal of several (G)RGG motifs. The mechanism by which RGG is reported to interact with RNA has been described previously^[72]^ and is unspecific to RNA sequence, with the exception of being able to recognize structural oligonucleotide motifs, such as G-quadruplexes.^[55,73-75]^

An RNA-sequence specific LLPS effect is more convincingly reported by the label at site 316, close to the carboxy-terminus in an S (30%)/ Y (10%) /R (17 %)-rich region. There are no RGG motifs in proximity to 316, yet the RNA sequence dependence is even stronger than at site 231. An immobile component (ii) can be clearly discerned in the presence of wt RNA and RNA*ga*, in addition to the broadened component (i), which is observed with all RNAs. Because the immobile component is well resolved at site 316, we can estimate that side-chain mobility at this site must change by at least an order of magnitude (*τ*_corr_(wt RNA) > 10 ns) between the free state and the state interacting with wt RNA (for additional spectral fitting see SI). Furthermore, the mobile component (m) is almost fully suppressed at site 316 in the sample with wt RNA.

For a possible explanation of the unexpected RNA sequence-dependent LLPS and mobility changes we consider the CW EPR observations in the context of the PRE measurements and of our ensemble model. For RNA-free, dispersed hnRNP A1, we had seen that the G-rich domain is mostly proximal to RNA binding sites in the RRMs (e.g. V90). This is illustrated by mapping the RNA from the 1:1 model by Beusch et al.^[51]^ onto UP1 color-coded by PRE intensity for spin labelling site 231 (same color-code as Figure 2(B)). The sites of strong PRE on the RRMs overlap with sites that recognize nucleobases in the RNA (Figure 4 (G)). The aromatic and H-bonding residues in the G-rich domain may thus act as intramolecular competitors for RNA binding. The mutated RNAs would then be expected to be less efficient at displacing the G-rich domain from the RRM surface, whereas the more strongly interacting wild-type motif would efficiently displace it. This would in turn frustrate the binding preferences of the G-rich domain and expose it to an aqueous environment. After such displacement from the RRM sites, the G-rich domain is positioned close to the RRM-bound RNA, possibly facilitating additional direct interactions, which we speculate might increase specificity in the binding Figure 4 (F). This may explain the strong effect at site 316, which is surrounded by a large fraction of amino acids with hydrogen bonding capacity.

Such selectivity due to cooperative binding of the RRMs and the G-rich domain to RNA could further act as a possible regulatory mechanism. Weak differences in stability of binding of the RNA to the RRM surface might thus be enhanced by additional interaction with the G-rich domain. Target sequences could thus be more tightly bound and more strongly immobilized than similar non-target sequences, limiting promiscuity of hnRNP A1 in RNA binding. In such a scenario, additional regulation might be achieved by post-translational modification of sites in the G-rich domain. Phosphorylation of serine, a major regulator of hnRNP A1 cellular localization, ^[6]^ introduces local negative charge in a previously uncharged region, and changes hydrogen bonding preferences, which might trigger the release of the bound RNA cargo after successful transport.

Finally, our ensemble model implies displacement of the G-rich domain from the RRMs upon RNA binding. This displacement could explain the importance of the G-rich region in hnRNP A1 multimerization^[25]^ and for LLPS. The unsatisfied preferences of this domain for contact with protein sites are expected to favor LLPS, which is indeed enhanced most strongly by specifically interacting with RNA.

## Conclusion

We report that specific RNA-binding by hnRNP A1 is tightly linked to LLPS. Already sub-stochiometric addition of specific RNAs, which carry two, one or no AG dinucleotide motifs embedded in the natural SMN2-ISS-N1 target sequence, cause such LLPS. While at high protein concentrations (100 μM) LDs appear with any of the tested RNA variants, RNA-sequence dependent LLPS was observed at lower protein concentrations (25 μM). DEER experiments on singly-labeled hnRNP A1 as well as SAXS and intermolecular PRE experiments revealed a minor fraction of oligomeric hnRNP A1 even at low protein concentration and in the absence of RNA. Both PRE data and an ensemble model of full-length hnRNP A1 revealed a compact conformation of the G-rich domain and its interaction with the RNA-binding face of the RRMs in the dispersed state. CW EPR experiments on singly spin-labelled mutants detected lower sidechain mobility in the presence of RNA. This effect is slightly more prominent with the wild-type RNA than with the mutated RNA*aa* and depends on the placement of the spin label in the G-rich domain. Our findings demonstrate that binding of specific target ssRNAs to hnRNP A1 can trigger phase separation and that single-point mutations in such RNA can significantly impact this process. We speculate that other proteins consisting of coupled RNA-binding and disordered domains may also exhibit such behavior, as it would provide some specificity to the complex interaction networks of phase-separating proteins that are promiscuous RNA binders.

## Acknowledgements

We would like to thank Emil Dedic and Georg Dorn for help with the SAXS acquisition and data processing. The ensemble SAXS experiments were performed with an instrument that was kindly provided by the Mezzenga lab at ETH Zurich. We also want to thank Irene Beusch, Fred Damberger, Laura Esteban-Hofer, Christoph Gmeiner, and Daniel Klose for useful discussions. We are grateful to Tanja Mittag and co-workers for discussions about the PRE experiments.

