## Supplementary Information for "Phase separation of hnRNP A1 upon specific RNA-binding observed by magnetic resonance"

### 1. Additional experimental details

#### a. Cloning, protein expression and purification

The coding sequence of hnRNP A1 (ROA1_HUMAN, isoform A1-A, Uniprot identifier P09651-2, see Figure S1(A)) was amplified from pET-9d-hnRNP A1[1] by PCR with upstream BamHI and downstream XhoI restriction sites and subcloned between the corresponding sites of a modified pET-28a(+) based DNA plasmid[2]. The resulting construct encodes TEV cleavable N-terminally $His_6$-tagged hnRNP A1. The plasmid was transformed into *E.coli* BL21-CodonPlus (DE3)-RIL cells for expression. Mutants of full-length hnRNP A1 with only one single cysteine for spin labelling were generated using partial mismatch PCR (quick-change method) with adapted primers. The native cysteines in the RRMs of hnRNAP1 were mutated to serine (C43S) and alanine (C175A), because undesired attachment of spin label to these intermediately buried sites was found to lead to severe protein precipitation. Mutation of site 175 was critical, and several different constructs were tested (C175S (purification failed), C175A, C175T, C175V), of which C175A was chosen because it is found in the mouse homologue of hnRNP A1 (Uniprot: P49312). Equivalent Cys mutants for UP1 (hnRNP A1 residues 1-196) were made by deletion of the C-terminal domain from the hnRNP A1 plasmid using specifically designed overlapping primer PCR (list see below). HnRNP A1 and UP1 constructs (wild-type, single cysteine + background, double cysteine +background) for spin labelling were over-expressed in *E.coli* cell cultures. Cells cultures were grown to an optical density at 600 nm (OD600) of approximately 0.8 at 37°C, at which point the temperature was lowered to 30°C, and over-expression was induced by addition of 0.5 mM isopropyl-β-D-thiogalactopyranosid (IPTG). Cells were harvested by centrifugation approximately 16 h after induction, resuspended in lysis buffer (20 mM Tris, pH 8.0, 1M NaCl, 1 mM dithiothreitol (DTT), cOmplete™protease inhibitor cocktail tablet (Roche)) and lysed by passing three times through a microfluidizer (Instrumat AG, Switzerland). His-tagged hnRNP A1 was purified with Ni-NTA affinity chromatography on an AKTAprime plus FPLC (GE Healthcare). After the first pass and elution from the column, the $His_6$ tag was cleaved by recombinant $His_6$-tagged Tobacco Etch Virus (TEV) protease digest at an engineered amino terminal cleavage site, which results in a final protein construct with an additional glycine at position -1, as well as the point mutation M1G. After desalting on a HighPrep 26/10 desalting column (GE Healthcare), the tagged protease as well as the cleaved tag were removed by a second pass over the affinity chromatography column. Proteins were transferred to dispersion buffer (50 mM sodium phosphate, pH 6.5, 100 mM L-arginine, 100 mM L-glutamate) by gravity flow desalting columns, and flash frozen in liquid nitrogen for storage at -80°C. Purity was confirmed by Coomassie stained SDS-PAGE (Figure S1(B)).

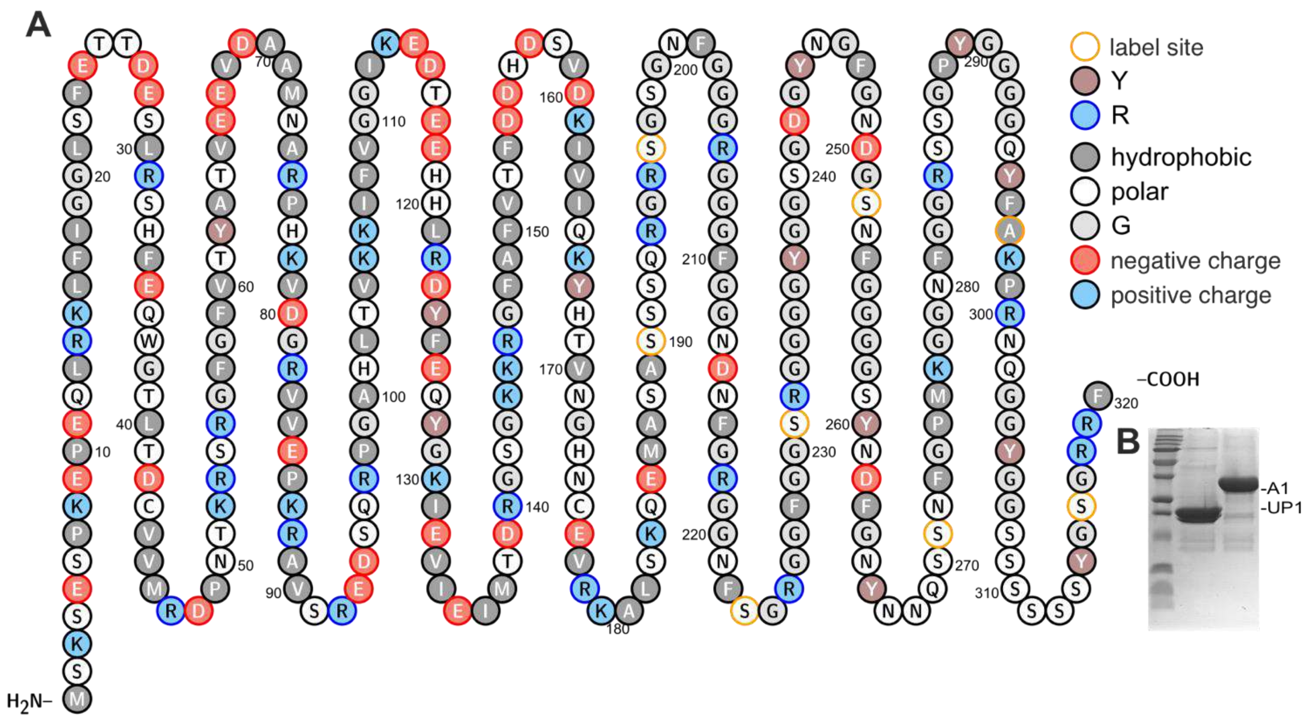


*Figure S 1* (A) Schematic representation of hnRNP A1. The IDD (in this work defined as 188-320) can easily be recognized by the high fraction of glycine residues. (B) Coomassie-stained SDS-Page of purified UP1 (middle) and hnRNP A1 (right)

**b.** Spin labelling

Cys-mutants designated for spin labelling were incubated at reducing conditions (2 h with 5 mM DTT at ambient temperature), and the protein was rebuffered to dispersion buffer which was pre-loaded with 10 x molar excess of S-(1-oxyl-2,2,5,5-tetramethyl-2,5-dihydro-1H-pyrrol-3-yl)methyl methanesulfonothioate (MTSL) for spin labelling. Spin labelling was performed at a low protein concentration (5-10 µM) to suppress the undesired side-reaction of protein dimerization via the solvent-exposed, reactive engineered Cys. At these conditions no enhanced hnRNP A1 dimer bands were observed in non-reducing vs reducing conditions for SDS-PAGE. After up to 16 h incubation at ambient temperature (no shaking), the residual free spin label was removed by gravity flow desalting columns (PD-10

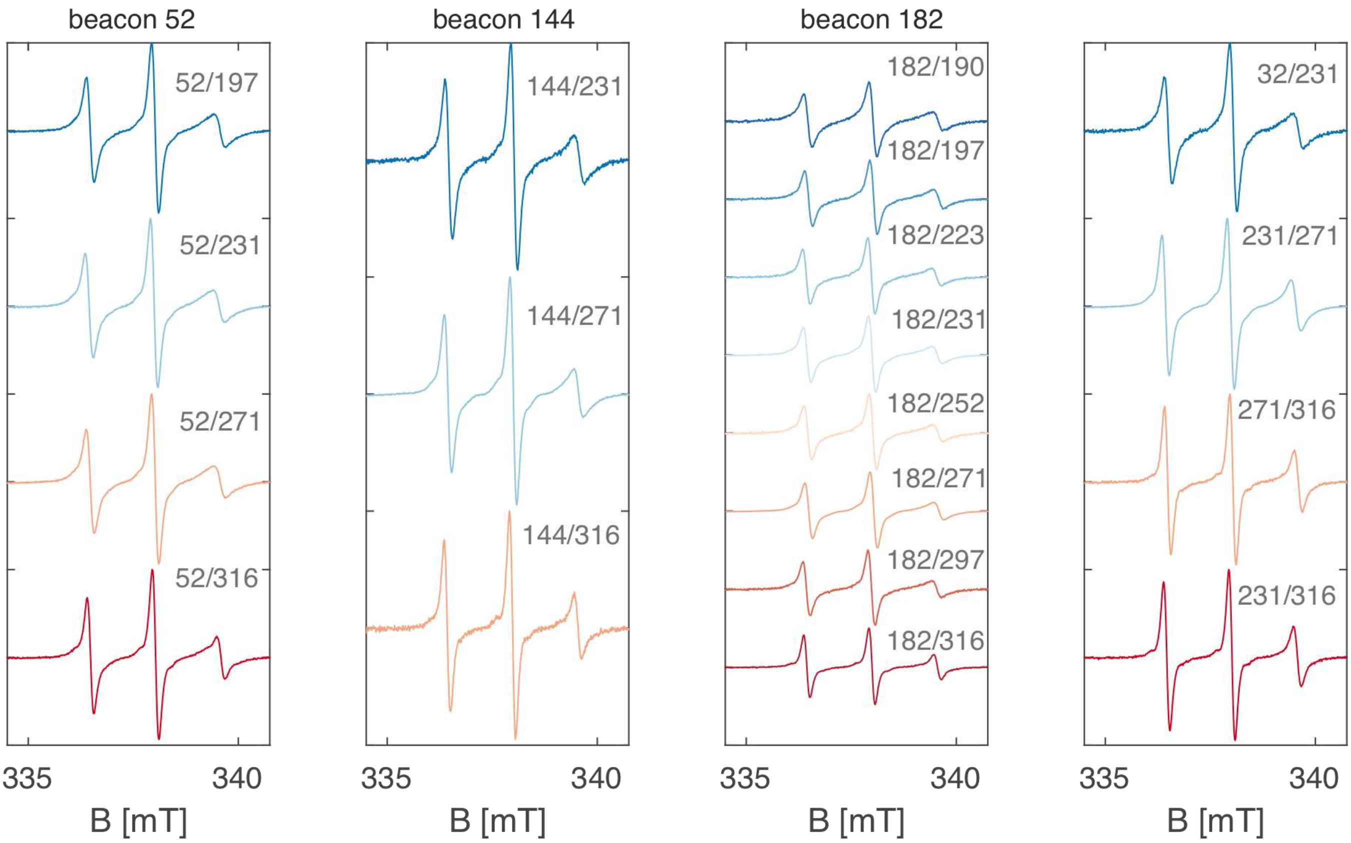


*Figure S 2* CW EPR X-band spectra of double Cys mutants of hnRNP A1 labelled with MTSL used in this study.

columns, or equivalent G-25 Mini-, resp. Midi-trap columns, depending on sample volume, GE-Healthcare). The labelling efficiency was estimated by spin counting with X-band CW EPR spectroscopy using a 100 µM 2,2,6,6-tetramethylpiperidin-1-yl)oxyl (TEMPO) reference solution. Spin labelled protein solutions were concentrated to approximately 50 µM in Amicon centricons (10 kDa MWCO, Merck), aliquoted, and flash-frozen by immersion in liquid nitrogen. We found that spin labelled hnRNP A1 should be frozen at concentration not exceeding 50 µM in order to have controlled, reproducible starting conditions after thawing a protein batch. At higher concentrations, we occasionally observed aggregation after/during thawing. In our hands, spin labelling with a maleimide-based (non-cleavable) nitroxide spin label led to severe protein precipitation for several spin labelling sites and was thus not further pursued.

c. Continuous-wave (CW) X-band EPR spectroscopy

CW EPR X-band spectra for spin counting for single and double Cys mutants are shown in Figure S2 (spectra normalized to the intensity maximum). All CW X-band EPR experiments were performed at 23 dB attenuation (1 mW power) and ambient temperature with an ELEXSYS-II E500 CW-EPR spectrometer equipped with a SHQ cavity using manual tuning and matching. For labelling efficiency measurements, the spin labelled proteins were measured in dispersion buffer without agarose, and 20 µl sample were loaded into small-diameter quartz capillary tubes (Blaubrand), which had been flame-sealed at one end. The protein concentration of each sample was determined with the same sample solution as was used for CW EPR by UV extinction measurement on a Nanodrop (ThermoFisher). For the RNA-binding study, the spin labeled protein stocks (50 µM) were thawed and mixed 1:1 (v/v) with pre-warmed (37°C) buffer mixed with 0.2% (w/v) low-melting temperature agarose (analogous to the imaging experiments, for agarose buffer preparation see below) containing the respective (or no) ssRNA. Unless stated otherwise, the CW EPR experiments were thus performed at 25 µM protein concentration. 15 µl sample were transferred to a pre-warmed, non-sealed X-band capillary by immersing the open capillary tube into the mixed solution at 37°C and carefully inverting the Eppendorf cup to drain the complete sample volume into the capillary, which is facilitated by the capillary effect. Any small air bubbles that entered the tube in this filling process were left to rise to the top of the solution while the agarose buffer was still warm and liquid. Filled capillaries were then left to cool to 25°C, where the agarose gel set (1-2 minutes). After setting, the gel cannot drain out of the capillary any more, but capillaries were nonetheless sealed with parafilm at the sample end to prevent evaporation. This sealing did not affect spectrometer tuning or yield any artifact EPR signals.

d. DEER

20-35 µl sample were prepared in 3 mm outer diameters quartz capillaries (New Era) for the DEER experiments. For cryoprotection, we either used 50% (v/v) d8-glycerol, or 0.2% (w/v) low-melting agarose solution. Samples using agarose were filled into pre-warmed capillaries (37°C) to avoid gel setting during the sample filling. The samples in the intramolecular distance measurements series for model generation of dispersed, RNA-free hnRNP A1 were all made with glycerol as the cryo-protectant and were directly frozen in liquid nitrogen. The samples for the RNA-related studies were flash frozen in liquid-nitrogen pre-cooled iso-pentane to achieve a faster cooling rate, which reduced the risk of damage to the tube due to sample expansion. Conventional four-pulse DEER was measured at 50 K (set by a liquid He cryostat) on a homebuilt Q-band pulse EPR spectrometer in a home-built resonator for oversized samples[3]. All pulse lengths were set to 16 ns, and a frequency separation of 100 MHz was used between pump position (on the maximum of the nitroxide spectrum) and detection position. The first refocusing delay was set to 400 ns, and the second refocusing was set to the maximum delay that still gave a visible stationary DEER echo at the zero-time with 100 shots averaging. The DEER traces were averaged for 12 h to 48 h.

e. Paramagnetic relaxation enhancement (PRE) experiments

$^{15}$N-enriched mutants of Cys-free hnRNP A1 (C43S, C175A) and single-Cys mutants (C43S, C175A, S231C), (C43S, C175A, S271C), (C43S, C175A, S316C) were purified analogously to samples without isotope enrichment after growth of bacterial cultures in M9-minimal medium with $^{15}$N-ammonium nitrate (0.5 g/l) as the nitrogen source in the medium, according to previously reported protocols.[2] Natural abundance (practically NMR-silent) hnRNP A1 single Cys mutant for intermolecular PRE experiments was already available from previous CW X-band EPR experiments. Spin labelling of the paramagnetic samples was performed with MTSL under the same conditions as for CW EPR and DEER samples. For the reference spectrum, we used the same batch of isotope-labelled hnRNP A1 mutant, and performed labelling with a diamagnetic analogue of the spin label (acetylated MTSL, here 'R1ac', (1-acetoxy-2,2,5,5-tetramethyl-δ-3-pyrroline-3-methyl)methanethiosulfonate, CAS: 392718- 69-3, Toronto Research Chemicals, Canada). Labelling with R1ac was performed with the same protocol as for MTSL, except that attachment of 'R1ac' was confirmed not by CW EPR, but by electron spray ionization mass spectrometry (ESI-MS, performed by the bioanalytic service at University Zurich) which showed the expected mass shift. NMR measurements were performed on a Bruker 700 MHz NMR spectrometer equipped with an AVANCE NEO console and a CryoProbe at 40 µM $^{15}$N-hnRNP A1-S231R1ac concentration (reference spectrum) and (40µM $^{15}$N-hnRNP A1 + 20µM non-isotope enriched hnRNP A1-S231R1) concentration in dispersion buffer. The backbone resonances of the diamagnetic analogue spectrum were slightly different from the assignment of wild-type UP1 available in the Allain group. We therefore purified $^{13}$C-$^{15}$N-enriched UP1 Cys-free (C43S, C175) mutant using $^{13}$C-glucose (1 g/l) as the carbon source in the M9-minimal medium, and assigned the backbone of hnRNP A1 Cys-free based on triple-resonance NMR experiments (HNCA, HNCACB, HNCO, HN(CA)CO, HN(CO)CACB). The buffer concentration in the measurement with the inter-molecular PRE experiment was slightly lower (10 mM sodium phosphate), but chemical shift perturbations were found to be minimal. 3% $D_2O$ were added to all samples for spectrometer locking, and samples were measured in 5 mm (outer diameter, 5TA, ARMAR Chemicals) throw-away tubes. Data was recorded and processed using Topspin (Bruker). Data acquisition of the 2D-$^{15}$N-$^{1}$H-HSQC spectra was for approximately 12 h per sample.

f. SAXS

Samples of wild-type hnRNP A1 and UP1 were thawed from purified stocks in dispersion buffer. After thawing, the samples were spun for 20 minutes at 20k x rcf, 20°C to pellet any aggregation from the freezing and thawing. The solutions were adjusted to 1.7 mg/ml (hnRNP A1), resp. 4.1 mg/ml (UP1) with dispersion buffer, and 100 µl each were transferred to quartz capillaries and sealed vacuum tight with glue. SAXS was measured by Dr. Georg Dorn and Dr. Emil Dedic using the facilities kindly provided by the Mezzenga lab at ETH Zurich.

g. ssRNA preparation

The ssRNAs used in this study were bought from Dharmacon (USA) and were shipped with 3' protection. Deprotection was performed according to the manufacturer's protocol, and the deprotected RNAs were resuspended after lyophilized in $ddH_2O$ at 1 mM. A 500 µM solution of each RNA was made in dispersion buffer (50 mM sodium phosphate, pH 6.5, 100 mM R/E), which was used all further experiments.

| Name | Sequence (5' → 3') |
|---|---|
| wild-type (RNAgg) | CCAGCAUUAUGAAAGUGA |
| RNAg*a* | CCAGCAUUAUGAAAAUGA |
| RNA*a*g | CCAACAUUAUGAAAGUGA |
| RNA*aa* | CCAACAUUAUGAAAAUGA |

h. Confocal imaging and agarose buffer handling

Confocal images were recorded in transmission mode on a Leica SP8 STED confocal microscope (488 nm illumination wavelength). The buffer for imaging was supplied with 0.2% (w/v) low-melting agarose to slow down droplet fusion and sinking in the LLPS state. Agarose solutions were prepared by weighing appropriate amounts of low-melting agarose powder in 2 ml Eppendorf tubes and mixing with dispersion buffer. To dissolve the agarose completely, the tubes were set to 95°C for about 10 minutes (until no more agarose clumps were visible). We found that the thus dissolved agarose solutions could be stored at 37°C in closed Eppendorf tubes (no shaking) for several days without setting of the gel. In contrast, the gel sets within a few minutes when placed at ambient temperature, and needs to be heated to 95°C again for melting. To make the imaging samples of hnRNP A1 (stored in dispersion buffer), we mixed the protein stock at the desired protein concentration in individual Eppendorf cups with dispersion buffer. For the experiments with RNA, an appropriate volume of RNA in dispersion buffer was pre-mixed with low-melting agarose-containing buffer and left at 37°C for one minute. The protein solution was added to the RNA-containing buffer, mixed briefly on a vortex, spun down briefly (< 1000 x rcf), and a 5 µl drop of sample was transferred to a chamber of an uncoated µ-well slide (ibidi, Germany). The lid was closed during drying, and the agarose gel set within less than a minute. Some buffer evaporation was always observed, which might slightly affect the final concentration, but the drops did not appear to shrink significantly in size. Pre-warming the glass cover plates was tested for slower gel setting, but it was found to strongly enhance buffer evaporation (and thus uncontrolled sample volume loss) before solidification of the agarose, and was thus not further pursued. The imaging chambers were sealed during measurement to prevent evaporation, but typically samples could not be stored for more than 12 h due to slow evaporation of water. For each condition, several drops were imaged in this manner. Unless stated otherwise the imaging was performed with wild-type hnRNP A1 resp. UP1. Images were recorded as 41x41 µm, unless stated otherwise, using a NA 1.2 water immersion objective.

i. Fluorescent imaging

Two types of fluorescent imaging experiments were performed. For detecting hnRNP A1 in the droplet, we used unspecifically surface labelled hnRNP A1. The fluorescent labelled hnRNP A1 was obtained by following the protocol for fluorescent protein labelling with the Monolith NT Protein Labeling kit BLUE-NHS (Nanotemper technologies, Munich). The solution was passed through a desalting spin column to remove excess dye. 0.5 µl labelled protein solution (unknown, but very low concentration) were used per 25 µl imaging sample solution. Fluorescent imaging was performed by excitation at 488 nm and detection above 510 nm. The second experiment for detecting RNA was performed by pre-mixing 0.25 µl GelRed Nucleic Acid stain (Biotium) with the hnRNP A1 solution prior to mixing with the RNA-agarose buffer solution. Fluorescent imaging was performed by excitation at 488 nm and detection above 510 nm.

j. Native PAGE

RNA-binding to the UP1 and hnRNP A1 constructs was confirmed by native PAGE (10% polyacrylamide gel (29:1), 0.5x TBE). Unless stated otherwise, 50 pmol RNA were loaded per lane, and UP1, resp. hnRNP A1 were added in the indicated molar ratio. The gels were run for 20 min at 120 V and stained with toluidine (RNA-stain). Destaining was performed in water until the desired contrast was reached.

k. Optical Density

Optical densities of the samples were measured in droplets buffer with additional 1.5% low melting agarose. For sample preparation 200 µM protein stock was mixed in 1:1 with 2 x agarose droplet buffer with corresponding RNA. Sample volume was 20 µl and pipetted into 384-well plate (Corning 4581) at 37°C while agarose still liquid. The small sample volume allowed temperature equilibration to 25°C within 10 minutes. Optical density was measured at 600nm using a Synergy 2 (Biotek) microplate reader. All measurements were experimental triplicates and the agarose buffer was used as a blank.

### 2. Data analysis procedures and additional spectroscopic data

#### a. Inter-beacon DEER and comparison between UP1 and hnRNP A1

DEER control experiments between the main beacon sites in the folded domains of hnRNP A1 (sites 52, 144, 182) were performed to confirm the validity of the solution state model of the RRMs as a starting point for the modelling of full-length hnRNP A1. The experimental data and fits with DeerAnalysis are shown in Figure S3. The zero-time for fitting was set to 120 ns. The data were first background corrected using a homogenous background model (dimensionality $n_{BG}$=3), and fitted with Tikhonov-regularization using the L-curve criterion for selection of optimal regularization parameter. Validation was performed using the default background start and white noise options. For two of the sites, we also performed analogous measurements with UP1 (lacking C-terminal residues 197-320 w.r..t full length hnRNP A1). As expected, the absence of this disordered domain had only minimal effects on the inter-beacon distance measurements. The simulation curves in Figure S3 were obtained with the DEER module in MMM,[4] by *in silico* labelling of the respective sites in the solution model of UP1 (PBB: 2LYV[2]) with the standard MTSL rotamer library at 298K.

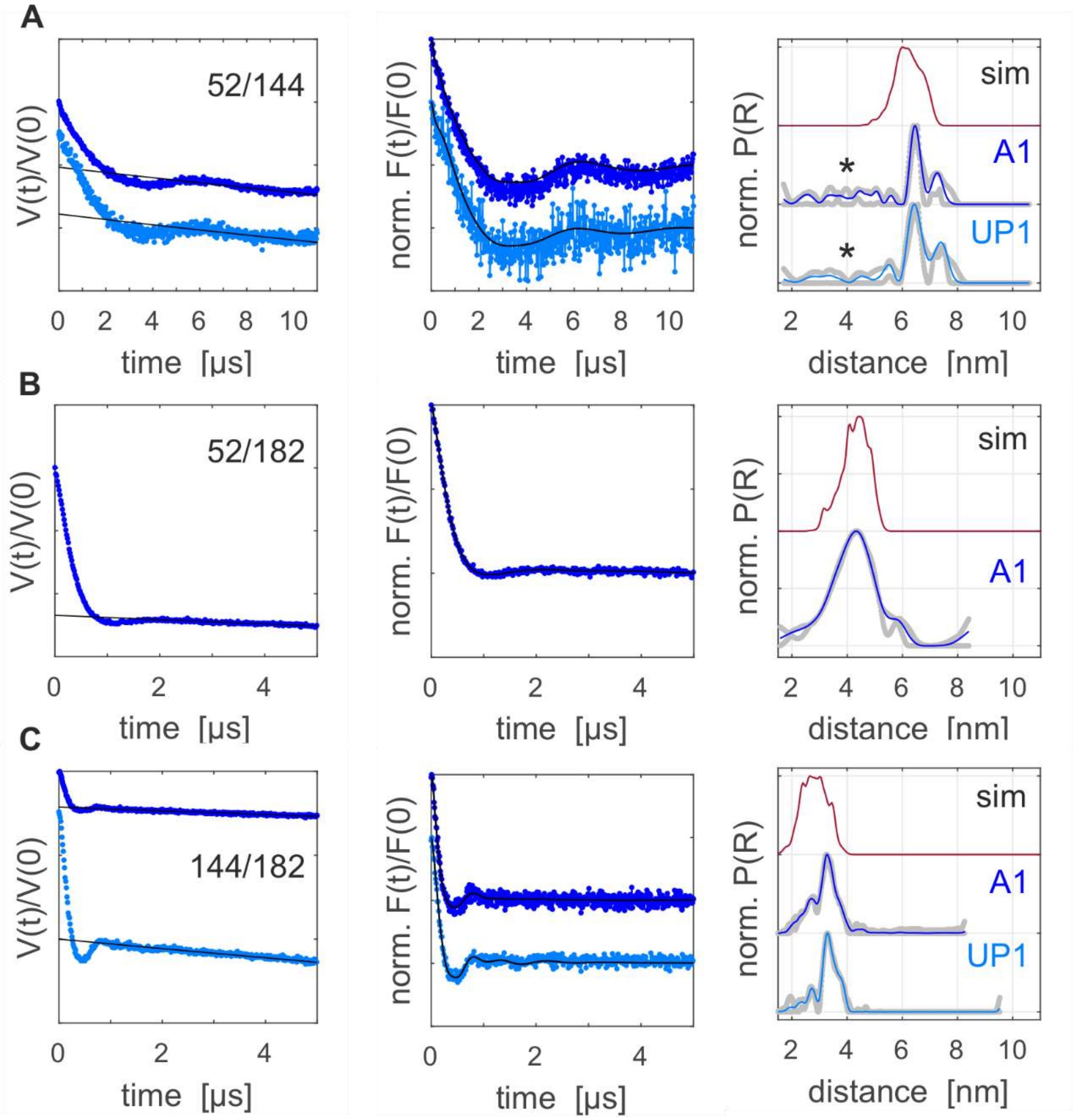


*Figure S 3* Inter-beacon DEER measurements: primary data V(t)/V(0), form factors F(t)/F(0) and model-free Tikhonov fitted distance distributions P(R) with validation error limits between beacon sites (A) 52/144; (B) 52/182; (C) 144/182 in the folded domains of hnRNP A1 as measured with the full length protein construct (1-320, ‘A1’), resp. in the RRM-only construct (1-196, ‘UP1’). MMM simulations with the solution structure of UP1 (pdb: 2LYV) are shown in red (MTSL rotamers at 298K);

b. DEER for intramolecular distance distribution measurement as ensemble restraints

Due to the more convenient simultaneous handling of multiple datasets, the DEER data for the ensemble modelling restraints were fitted with DeerLab (https://jeschkelab.github.io/DeerLab).The routine automatically detected experimental zero-time (~120 ns), then data were background corrected using a homogenous background model (dimensionality $n_{BG}$=3). The resulting form factors were then fitted by a single Gaussian with variable mean distance *r* and variable distribution width *σ* (standard deviation), unless stated otherwise. In Figure S4 we show the primary data, fits (red), and background contribution of the fit (grey extrapolated line) for all available 19 DEER distance restraints. The resulting distance distributions and error estimation ranges using a grid-based screening of the modulation depth Δ, white noise parameter and background fit range parameter (analogous to using 'DeerAnalysis 'validation' tool) are shown in Figure S5. The Gaussian fit parameters are reported in Table S1. The labelling efficiency η= c(spin)/c(Cys)= c(spin)/ [$n_{Cys}$*c(protein)] as determined by spin counting with CW EPR spectroscopy (see CW EPR section below) is included in the same table.

| Site A | Site B | R [Å] | σ [Å] | η [%] |
|---|---|---|---|---|
| 32 | 231 | 51.2 | 15.5 | 74 |
| 52 | 197 | 35.6 | 13.8 | 73 |
| 52 | 231 | 37.5 | 18.1 | 93 |
| 52 | 271 | 28.1 | 19.7 | 84 |
| 52 | 316 | 37.5 | 18.2 | 77 |
| 144 | 231 | 49.7 | 15.4 | 44 |
| 144 | 271 | 45.7 | 18.6 | 70 |
| 144 | 316 | 44.1 | 11.7 | 55 |
| 182 | 190 | 22.5 | 6.2 | 51 |
| 182 | 197 | 27.4 | 9.5 | 65 |
| 182 | 223 | 31.4 | 14.2 | 52 |
| 182 | 231 | 33.3 | 14.8 | 82 |
| 182 | 252 | 36.6 | 12.7 | 67 |
| 182 | 271 | 30.5 | 18.8 | 73 |
| 182 | 297 | 34.1 | 15.8 | 54 |
| 182 | 316 | 34.8 | 15.7 | 68 |
| 231 | 271 | 20.1 | 15.3 | 68 |
| 231 | 316 | 27.1 | 16.5 | 72 |
| 271 | 316 | 22.6 | 15.8 | 48 |

*Table S 1* Intramolecular DEER fit parameters

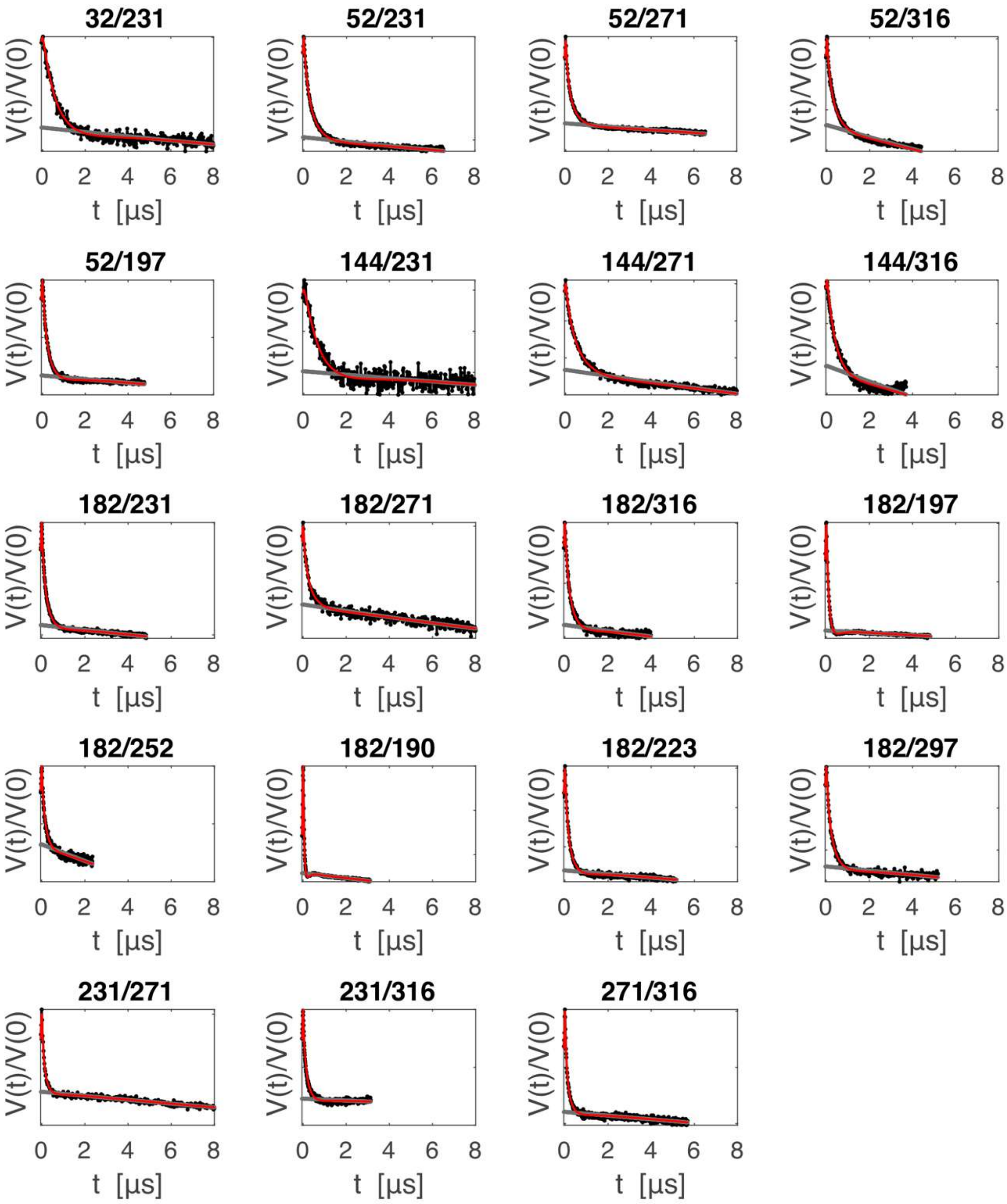


*Figure S 4* All 19 available DEER experiments; primary data V(t)/V(0) (black), homogenous background fit (grey) and single-Gaussian fit (red, combined background and form factor contribution);

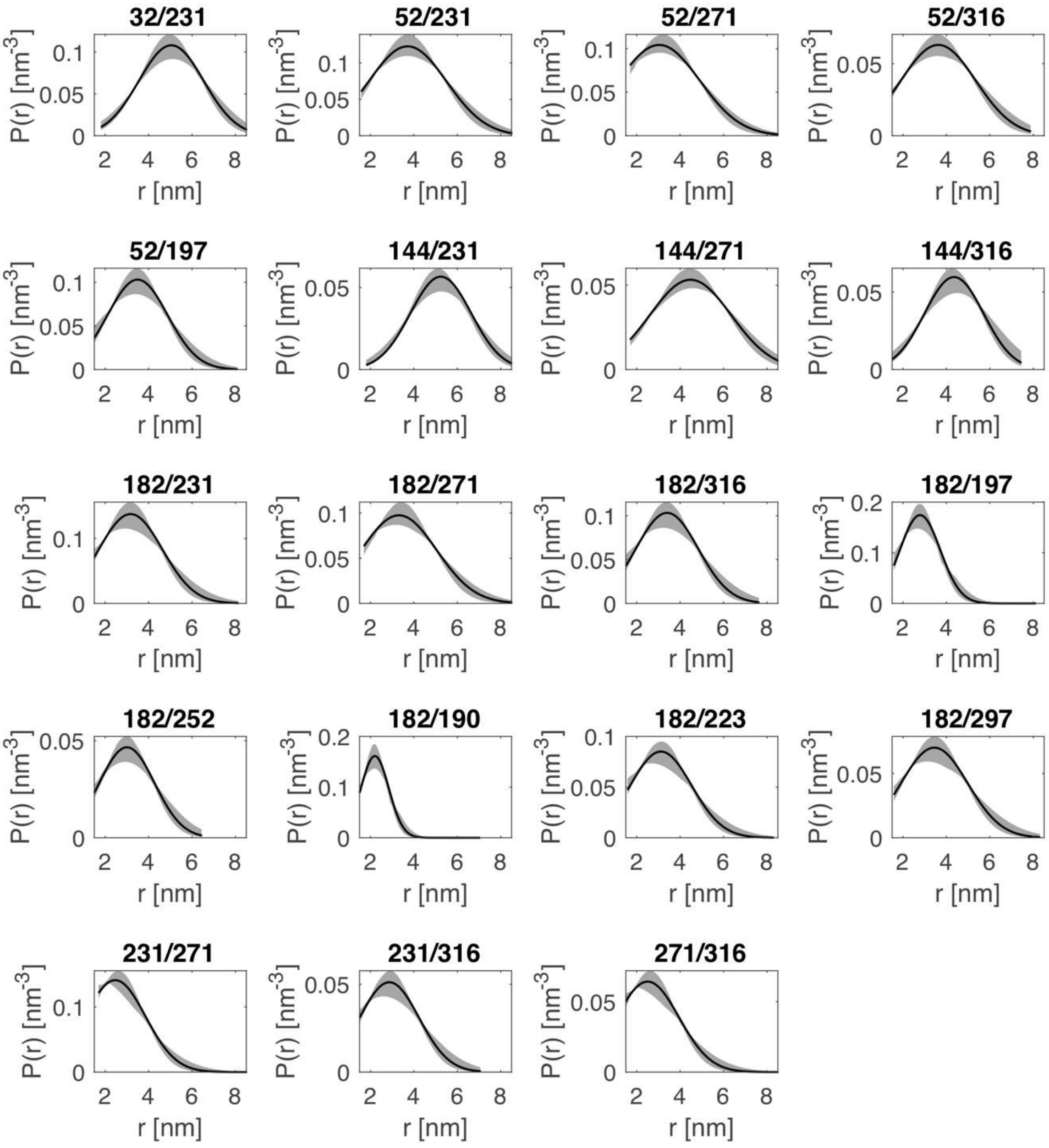


*Figure S 5* Fitted single Gaussian distance distributions (black) for data in Figure S4 and error estimation ranges (grey areas) from validation-type analysis;

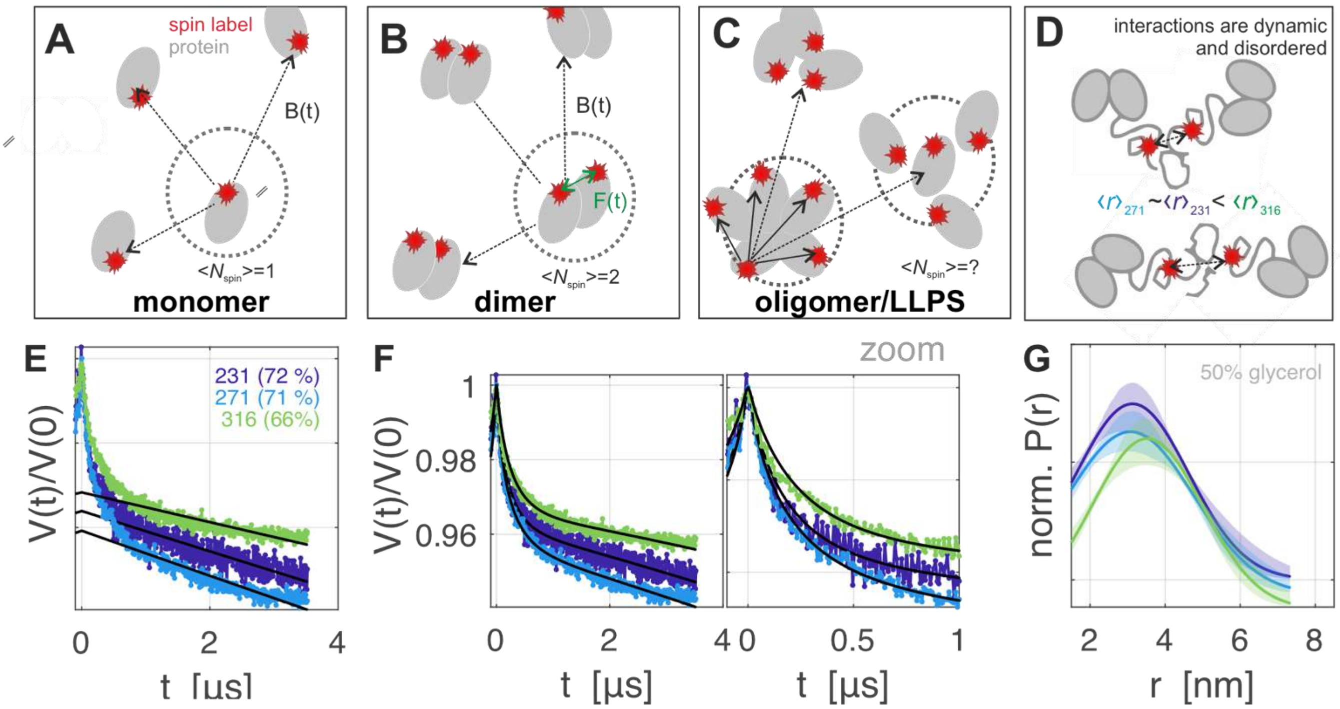


*Figure S 6* (A-D) schematic representation of how protein-protein interactions can lead to DEER modulation (A) dispersed singly labelled monomer (average spin per object $\langle N_{spin}\rangle=1$) gives only background signal B(t). (B) dispersed doubly labelled dimers (average spin per object $\langle N_{spin}\rangle=2$) give an additional form factor contribution F(t); (C) oligomeric/phase separated singly labelled proteins form a complicated intermediate case with ill-defined $\langle N_{spin}\rangle$, and different spin relaxation rates between dense and dilute phase. (D) interactions between disordered domains are also disordered (E-G) distance distribution analysis of DEER measured with singly spin labeled hnRNP A1 at three sites in the G-rich domain, which shows a minor contribution from interacting molecules as steep initial decays.

c. Intermolecular DEER analysis of free hnRNP A1

DEER with singly spin-labelled proteins was applied to characterize protein-protein interactions. Contributions to the DEER signal that enable this analysis are schematically displayed in Figure S6. The DEER primary data for hnRNP A1 singly spin-labelled at sites 231, 271, and 316 are shown in Figure S6, analogous to the data in main text Figure 1(H). The samples in this series were at 50 µM protein concentration, labelling efficiency as determined by spin counting is annotated (no spin dilution). Deuterated glycerol 50% (v:v) was used as cryoprotectant in this series. A distance analysis was performed in the same way as for the distance restraints (homogenous background correction followed by single-Gaussian fit) and is shown as well. This analysis implicitly assumes that all of the short-range interactions that are not removed by the background fit arise from dimerized protein and that the monomers and dimers are otherwise homogenously distributed in the sample (background dimensionality of the DEER fit is three). The distance distributions are very broad and very similar for all three sites, which demonstrates that the proteins are not interacting in a well-defined manner, but rather form a disordered complex or a condensed phase. Note that distance data derived in this way can only serve as a guideline to monitor protein-protein interactions. They are probably a poor approximation of the actual distance distributions, given that a fraction of hnRNP A1 is likely in the LD state, where local spin label concentration is expected to be very high.

d. Intermolecular DEER with hnRNP A1 in the presence of RNA

DEER was also measured with singly spin-labelled hnRNP A1 in the presence of RNA. Since this is addition is known to induce LLPS at our conditions, we expected strongly enhanced background decay and more intermolecular interactions compared to free hnRNP A1, which was indeed observed, see Figure S7. We show DEER data for the three sites 52, 144 and 231 with two different cryoprotectant conditions: 50% (v:v) deuterated glycerol, resp. 1% agarose. As with the free hnRNP A1, we show distance analysis (homogenous background correction followed by Single Gaussian fit) as a guideline for interpreting differences in the featureless decays. In all glycerol conditions, wtRNA leads to a somewhat steeper initial decay and thus to shorter distance contributions than RNA*aa*. In the presence of agarose, the separation between long-distance background and short-distance form factor range becomes more difficult, especially at site 231.

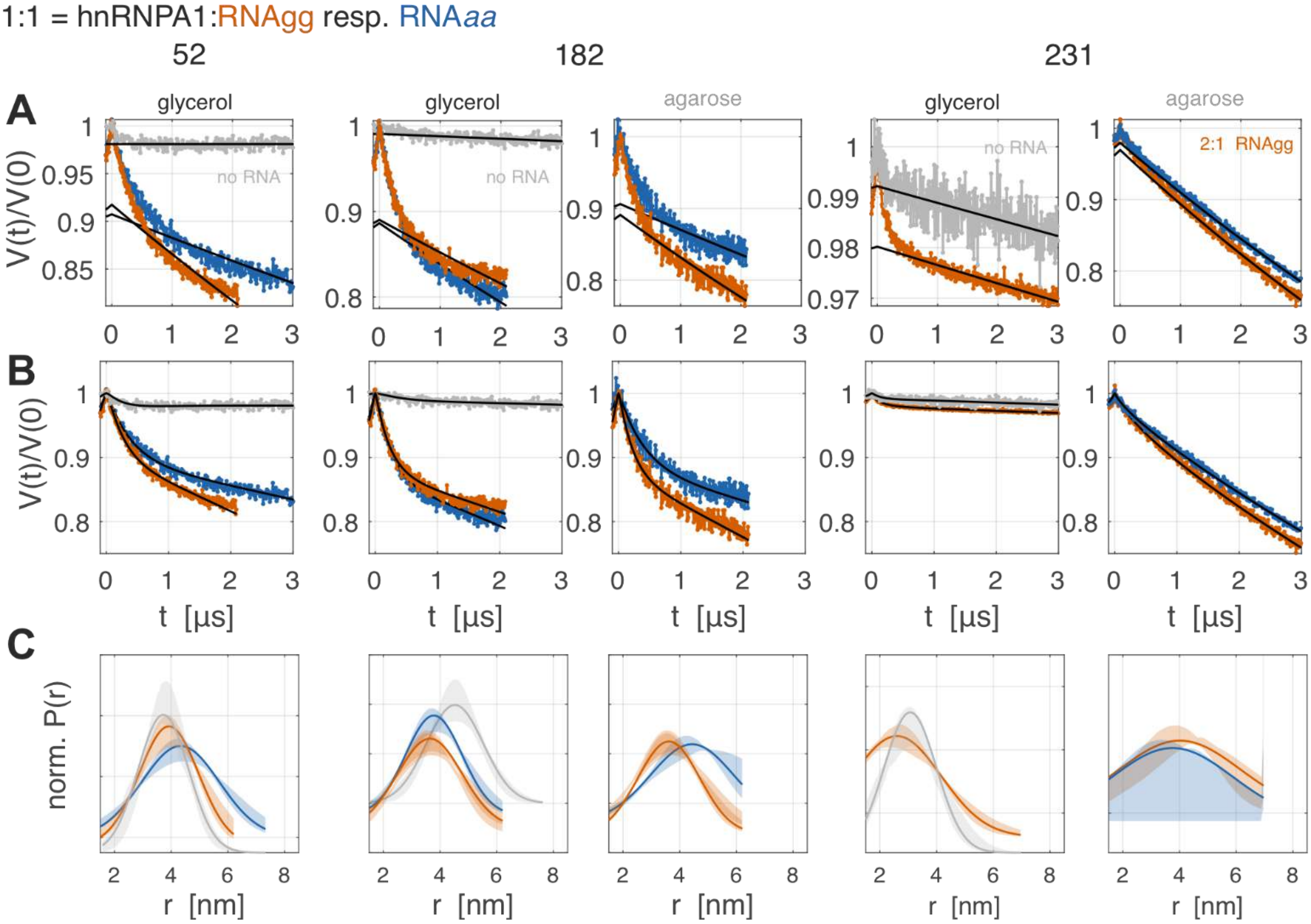


*Figure S 7* Intermolecular DEER at three different labeling sites in free hnRNP (grey) and in the presence of 1:1 molar ratio of wild-type RNA (RNAgg, orange) or RNA*aa* (blue).

e. <u>CW EPR analysis and fitting for spin labelling sites in the G-rich domain of hnRNP A1</u>

In the dispersed state we fitted the intermediate motion spectra obtained with the spin labels in the IDD of hnRNP A1 with an isotropic tumbling model using the ‘chili’ function in EasySpin.[6] The spectral parameters of the spin label MTSL in dispersion buffer were determined independently prior to fitting the CW-X band data of spin labelled hnRNP A1, to minimize the number of fit parameters in the latter case. To this end we fitted a spectrum of flash frozen, 100 µM MTSL in dispersion buffer at 140 K with the EasySpin ‘pepper’ function[6] to determine the anisotropic *g*-tensor parameters as well as the axial *A*-tensor parameters (see Figure S8). These were used to find the coupling frequency expected of four equivalent natural abundance $^{13}C$ satellites that could be observed in the spectrum obtained with 100 µM MTLS in the same buffer at ambient temperature with the ‘chili’ function with isotropic tumbling.

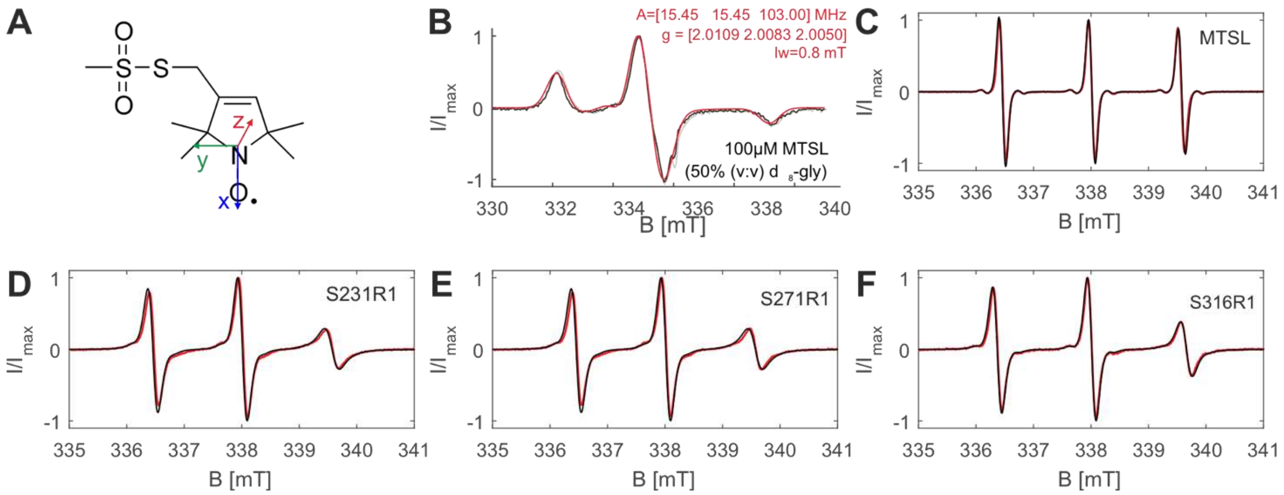


*Figure S8* Fitting of nitroxide spectra; (A) MTSL spin label (B) spectrum and EasySpin 'pepper' fit (red) of flash frozen solution of 100 μM MTSL in dispersion buffer at 140 K (C) spectrum and 'chili' fit (red) of 100 μM MTSL at ambient temperature using the spin Hamiltonian parameters from the fit in (B) and fitting only linewidth, isotropic tumbling rate $\tau_{corr}$,, and $^{13}$C isotropic hyperfine coupling of four equivalent carbons; (D-F) singly spin-labelled hnRNP A1 samples (no RNA) fitted using the spin system of free MTSL and adjusting only linewidth and $\tau_{corr}$,; spectra were collected a protein concentration of 25 μM and ambient temperature (no agarose stabilization);

The fitted free-label spin-system *g*-tensor and *A*-tensor parameters (*g* = [2.0109, 2.0083, 2.0050], *A*=[15.45, 15.45, 103.00] MHz, lw=0.8 mT) were used as the starting point for fitting the mutant spectra, by adjusting for each component only the linewidth (lw) and the value of the isotropic tumbling rate $\tau_c$. For the samples containing RNA, a single-component fit usually gave poor fit results, and consequently two- and three-component fits were performed, which introduced additionally the relative weights as fit parameters. To accelerate convergence, fitting was performed by four empirically derived stages of fitting on the Euler high-performance computing unit of ETH Zurich, with a total of up to 24 h fitting time per spectrum. The five fitting stages were: 1.) grid search with a rough grid (10% value change) optimizing fit parameters (lw, $\tau_c$, and weight per component); 2.) large-range (10%) simultaneous Levenberg-Marquard (LM) optimization of only one component weight and correlation time; 3.) large-range (10%) simultaneous LM optimization of all $\tau_c$ 4.) fine range (1%) repetition of stage 2; 5.) fine range (1%) LM fit of all fit parameters; the results are reported in Table S2 (two-component) and Table S3 (three-component. The overlays of fits and data are shown in Figure S9. Note that spin counting by double integration showed variations in total spin signal (Figure S10). Therefore, we cannot exclude that some, probably very broad spectral components escape this fit procedure. For those components that we are sensitive to, we clearly see that at least one component of strongly reduced mobility becomes dominant in the presence of RNA. The weight and correlation time are larger for wild-type RNA and RNAg*a* than for RNA*ag* and RNA*aa*. The equivalent spectral overlays and spin counting results for the RNA-concentration dependent series are shown in Figure S11. We quantified the relative spectral intensity in this titration at two resonance field positions corresponding to the low-field features of the fast resp. highly immobile components of type (ii), indicated by circles, resp. crosses in Figure S11(D,E)). The amplitude of the highly immobile component grows, as the mobile line decreases, following a sigmoidal shape that reaches a plateau after a 1:1 (RNA:protein) ratio for both RNAs. The plateau intensity ratio is approximately 1.5 times higher with the wt RNA, which most likely stems from a larger fraction of protein in the dense state. However, this ratio could be affected by how well the mobility in the dense state is described by a single rotational correlation time $\tau_{corr}$, as opposed to a distribution of $\tau_{corr}$.

**Table S2** Fitted spin-label mobility parameters for the two-component fit.

| Site [a] | RNA | 1000 x rmsd | 1000 x noise estimate | W1 [%] | $\tau_{corr}$ (1) [ns] | lw (1) [mT] | W2 [%] | $\tau_{corr}$ (2) [ns] | lw (2) [mT] |
|---|---|---|---|---|---|---|---|---|---|
| 231 | *aa* | 9 | 3 | 8 | 0.4 | 0.1 | 92 | 1.6 | 0 |
| | *ag* | 13 | 3 | 14 | 0.6 | 0.1 | 87 | 2.1 | 0.02 |
| | *ga* | 14 | 3 | 36 | 1.1 | 0.1 | 64 | 5.5 | 0.04 |
| | *gg* | 14 | 3 | 33 | 1.1 | 0.1 | 68 | 5.2 | 0.04 |
| | no | 12 | 5 | 81 | 0.8 | 0.1 | 19 | 11 | 0.03 |
| 271 | *aa* | 51 | 5 | 63 | 1.5 | 0.1 | 37 | 0.5 | 0.16 |
| | *ag* | 24 | 5 | 52 | 0.8 | 0.1 | 48 | 4.7 | 0.06 |
| | *ga* | 18 | 3 | 43 | 0.7 | 0.1 | 57 | 4.8 | 0.12 |
| | *gg* | 30 | 2 | 52 | 0.9 | 0.1 | 48 | 4.7 | 0.02 |
| | no | 13 | 6 | 79 | 0.7 | 0.1 | 21 | 9.9 | 0.04 |
| 316 | *aa* | 6 | 3 | 31 | 0.7 | 0.1 | 69 | 5.2 | 0.14 |
| | *ag* | 11 | 3 | 34 | 0.6 | 0.1 | 66 | 4.3 | 0.2 |
| | *ga* | 7 | 3 | 5 | 0.3 | 0.1 | 95 | 2.3 | 0.02 |
| | *gg* | 6 | 3 | 9 | 0.6 | 0.1 | 91 | 5.7 | 0.18 |
| | no | 37 | 6 | 70 | 0.6 | 0.1 | 30 | 0.3 | 0.17 |

**Table S3** Fitted spin-label mobility parameters for the three-component fit.

| Site [a] | RNA | 1000 x rmsd | 1000 x noise estimate | W1 [%] | $\tau_{corr}$ (1) [ns] | lw (1) [mT] | W2 [%] | $\tau_{corr}$ (2) [ns] | lw (2) [mT] | W3 [%] | $\tau_{corr}$ (3) [ns] | lw (3) [mT] |
|---|---|---|---|---|---|---|---|---|---|---|---|---|
| 231 | *aa* | 7 | 4 | 40 | 0.9 | 0.11 | 44 | 7.7 | 0.13 | 16 | 18.5 | 2.05 |
| | *ag* | 12 | 3 | 41 | 1.1 | 0.06 | 30 | 4 | 0.19 | 30 | 10.1 | 0.64 |
| | *ga* | 15 | 3 | 40 | 1.2 | 0.03 | 51 | 4.9 | 0.19 | 9 | 11.2 | 0.09 |
| | *gg* | 14 | 4 | 32 | 1.2 | -0.03 | 47 | 5.7 | 0.09 | 21 | 4.6 | 1.87 |
| | no | 10 | 5 | 67 | 0.6 | 0.14 | 11 | 3.6 | 0.15 | 22 | 6.9 | 0.85 |
| 271 | *aa* | 51 | 5 | 78 | 0.8 | 0.13 | 14 | 11.4 | 0.12 | 8 | 19.1 | 2.13 |
| | *ag* | 24 | 5 | 40 | 0.6 | 0.13 | 21 | 1.5 | 0.08 | 39 | 5.9 | 0.83 |
| | *ga* | 18 | 4 | 30 | 0.7 | 0.13 | 36 | 5.8 | 0.14 | 35 | 18 | 1.9 |
| | *gg* | 30 | 4 | 4 | 0.1 | 0.18 | 49 | 1.3 | 0.04 | 47 | 6.6 | 2.08 |
| | no | 12 | 6 | 74 | 0.7 | 0.12 | 6 | 4 | 0.15 | 19 | 7.9 | 0.85 |
| 316 | *aa* | 6 | 3 | 31 | 0.7 | 0.09 | 48 | 7.4 | 0.15 | 21 | 15.5 | 2.05 |
| | *ag* | 11 | 4 | 26 | 0.4 | 0.13 | 40 | 1.7 | 0.25 | 34 | 12.5 | 0.68 |
| | *ga* | 5 | 3 | 18 | 0.6 | 0.11 | 40 | 4 | 0.1 | 42 | 4.2 | 0 |
| | *gg* | 5 | 3 | 11 | 0.7 | 0.02 | 46 | 9.8 | 0.21 | 42 | 3 | 0.51 |
| | no | 38 | 6 | 71 | 0.4 | 0.13 | 7 | 8 | 0.2 | 22 | 17.5 | 1.13 |

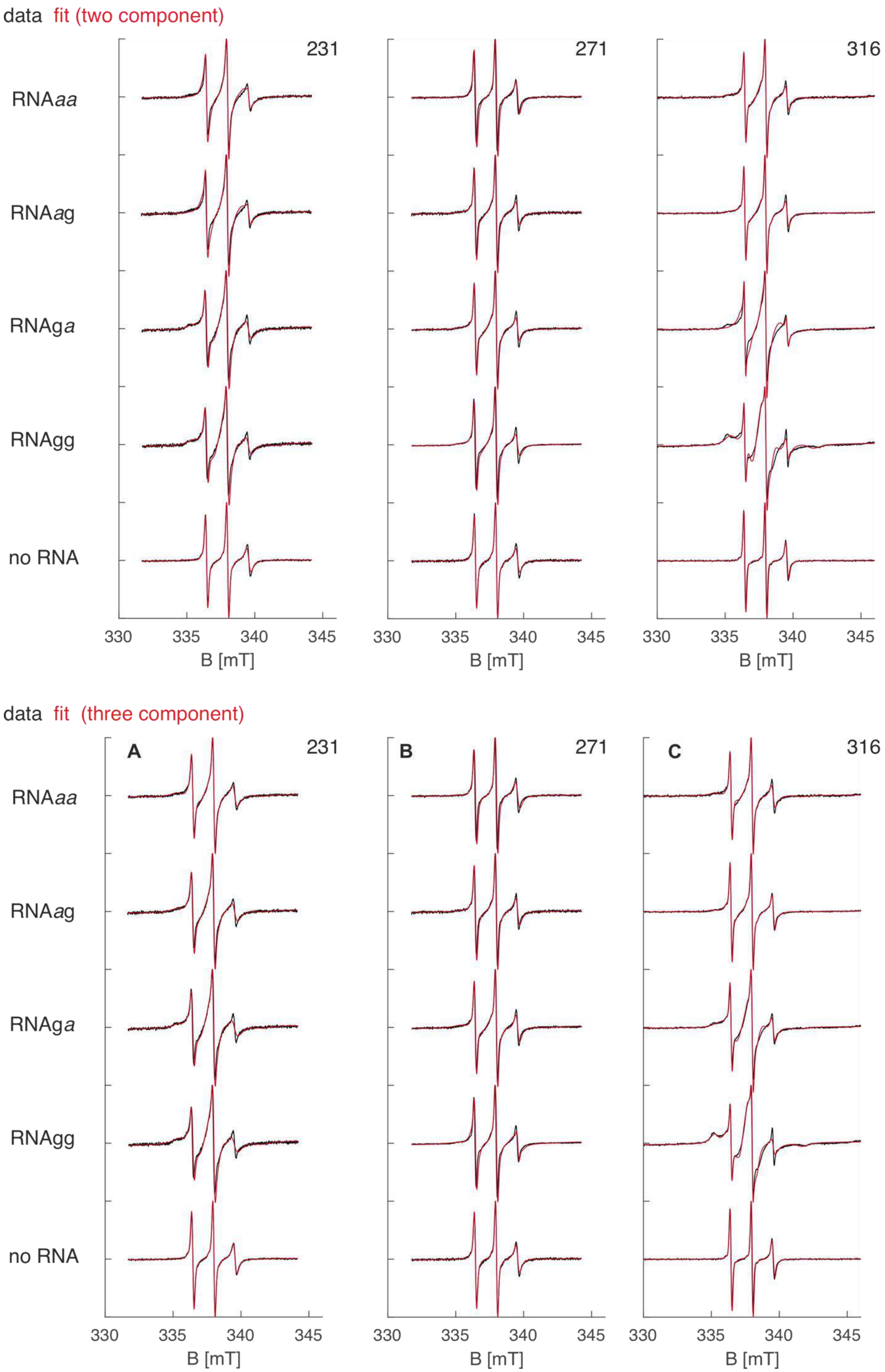


*Figure S9* CW EPR X-band spectra (black) of the singly spin labelled mutants 231, 271, and 316 of full length hnRNP A1 and two-component (top) as well as three-component (bottom) EasySpin ('chili') fits (red).

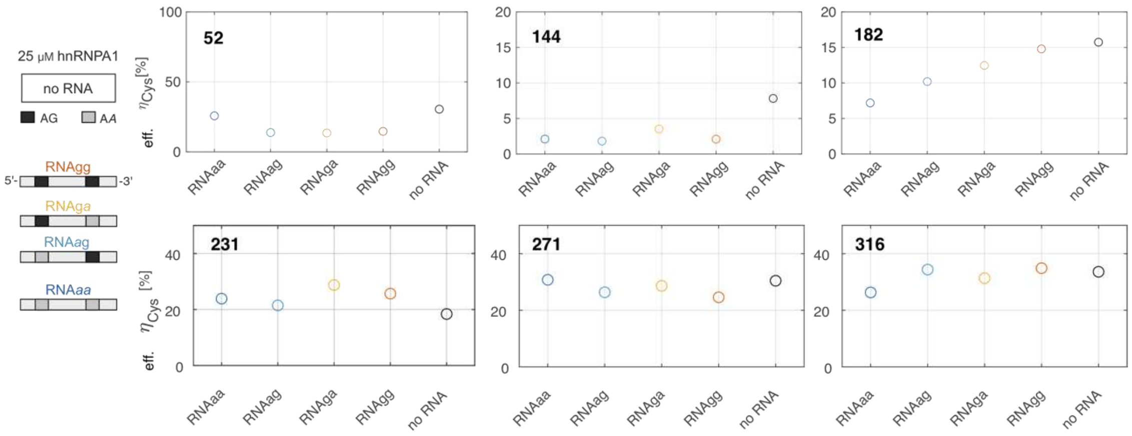


*Figure S 10* Spin counting for the CW EPR experiments with singly spin-labelled (sites indicated) free hnRNP A1 and hnRNP A1 + RNA (1:1 molar ratio). RNAs are color-coded as indicated on the left. The same batch of spin labelled-hnRNP A1 was used in each experiment. Hence, differences must arise from minor fluctuations in sensitivity, samples or from suppressed signal in the presence of RNA.

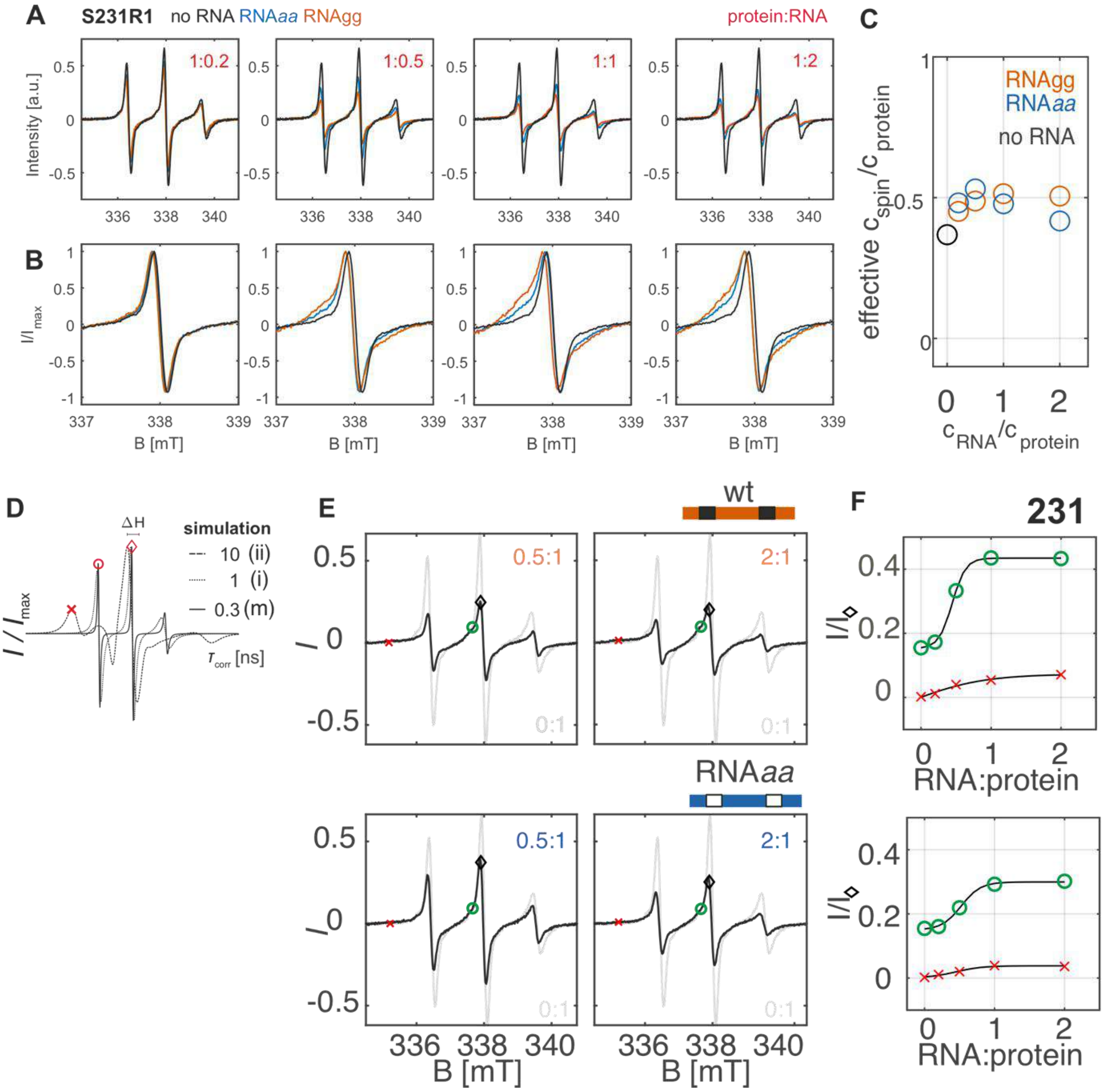


*Figure S 11* CW EPR X-band spectra of hnRNP A1 labelled at site 231 with increasing presence of RNA wt, and RNA*aa*; (A) unscaled full view; (B) scaled zoom on central transition; (C) effective spin counting in the series (RNAgg = wild-type RNA); (D) illustrative EasySpin simulation with different $\tau_{corr}$ ; (E) overlay an annotation of component analysis; (F) intensity fraction of components marked in (E) as a function of RNA:protein ratio;

a. <u>CW EPR analysis and fitting for spin labelling sites in folded domains of hnRNP A1</u>

The CW-EPR spectra of hnRNP A1 labelled at the beacon sites in the folded domains in the free state, as well as upon addition with 1:1 molar RNA variants in agarose stabilized conditions are shown in Figure S12. Overall, the effect of RNA binding appears weaker and unspecific to RNA sequence compared to the sites in the G-rich domain. The already broad and asymmetric appearance of the spectra in the free state compared to the labelling sites in the IDD introduces additional ill-defined fit parameters, and final fitting was thus not pursued.

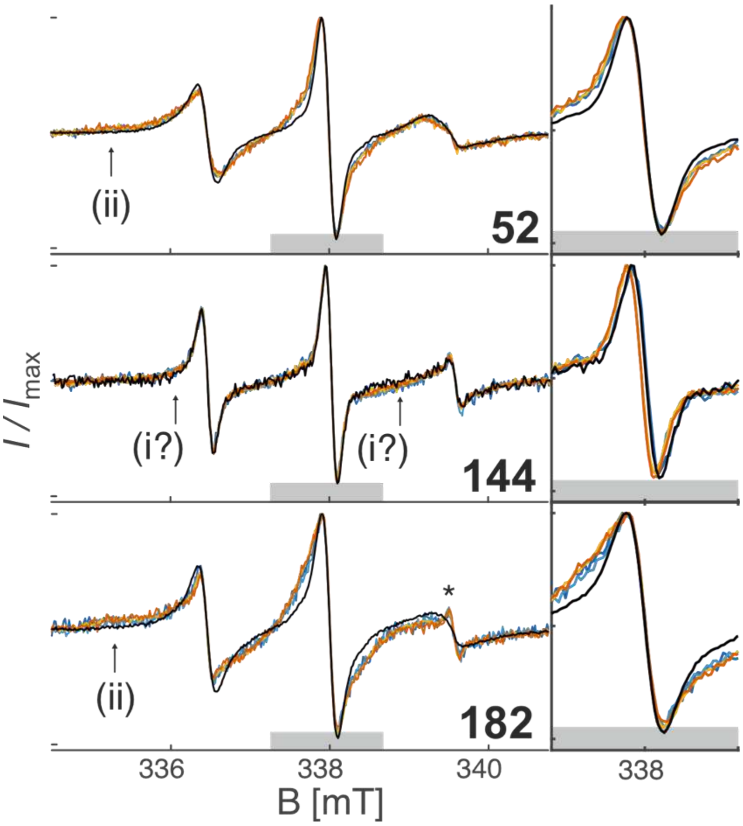


*Figure S 12* Normalized CW EPR X-band spectra and zoom to the central transition of hnRNP A1 singly spin-labelled at beacon sites in folded domains in free state (black) and in the presence of RNA variants (orange: wt RNA, yellow: RNAg*a*, light blue: RNA*a*g, dark blue: RNA*aa*); Some additional arising immobile (i) and very immobile (ii) components are labelled. The asterisk marks the appearance of a very minor, more flexible component at site 182;

3. <u>EPR results with UP1 in the presence of RNA</u>

Approximately 50 µM samples of MTSL-labelled double-Cys mutants of UP1 were incubated in a 1:1 molar ratio with wtRNA ('RNAgg') and in the series of mutated RNAs used in this study (RNA*ag* , RNA*ga* , RNA*aa*), mixed with 50% (v:v) d8-glycerol as cryo-protectant and flash frozen in liquid nitrogen-precooled iso-pentane. DEER experiments were recorded in the same way as for the distance-restraint measurements with free hnRNP A1 (see above). The data and DeerAnalysis Tikhonov regularization fits are shown in Figure S13. In contrast to hnRNP A1 in the presence of RNA, the background contribution remained low for UP1 in the presence of RNA, and no broadening was observed in ambient temperature CW EPR spectra. This is consistent with the observation that UP1 does not phase separate in the presence of RNA, as was confirmed by imaging (see imaging section below). The slight change in the distance distributions in UP1 upon adding RNA are broadly consistent with modelling based on the solution structure model by Beusch et al.,[5] as seen comparison with the simulations in Figure S13(D).

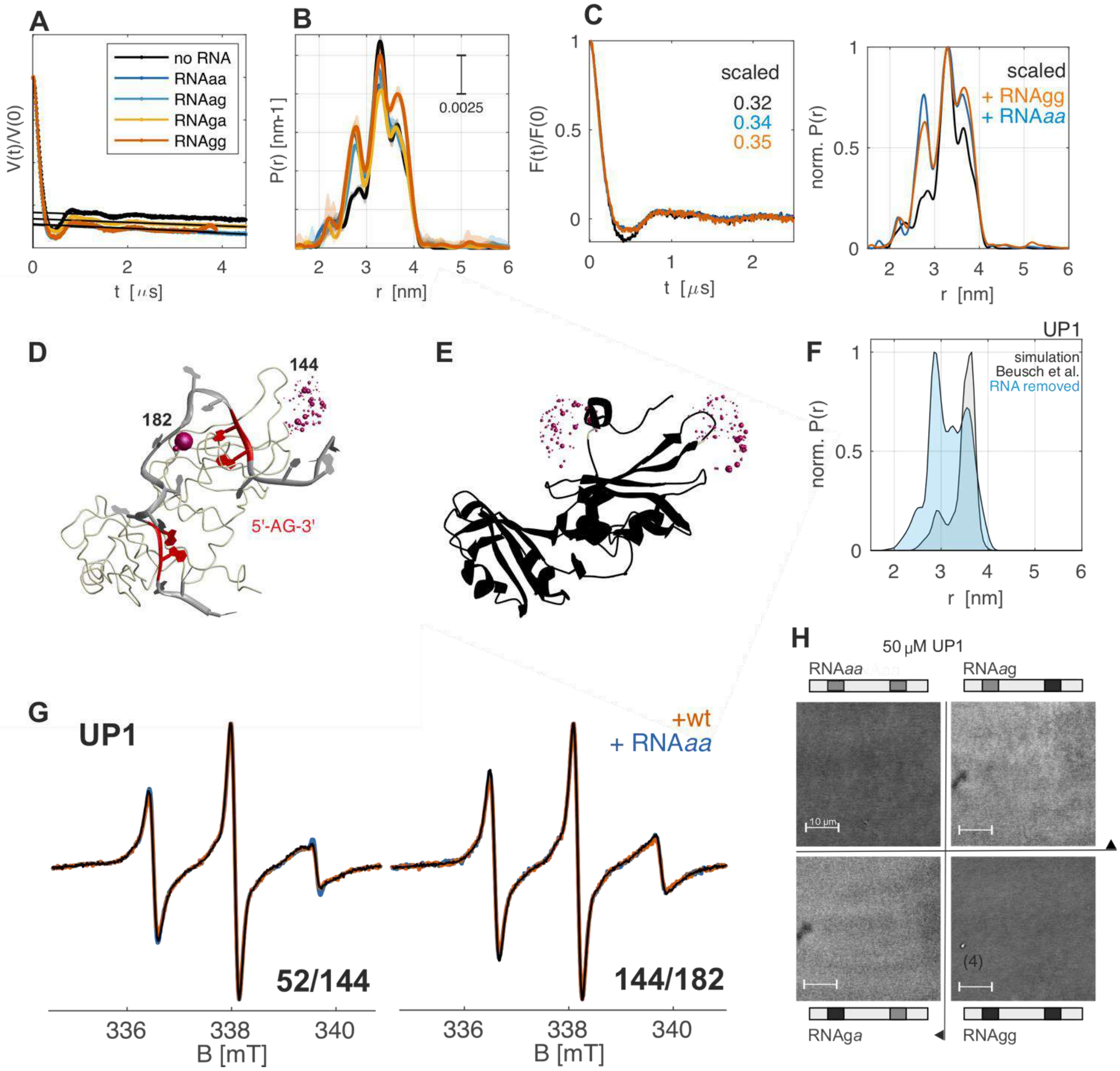


*Figure S 13* UP1 in the presence of RNA variants; (A) primary DEER data for UP1 MTSL-labelled at sites 144/182; free UP1 (black), and UP1 in the presence of 1:1 molar ratio RNA variants (color-coded); (B) fitted distance distributions (Tikhonov regularization); (C) modulation depth-scaled overlay of background corrected form-factors with free UP1 and UP1 incubated with wt RNA/RNA*aa* (left) and corresponding scaled distance distributions (right); (D) solution model of UP1 with highly similar RNA as used in this work (two 'AG' motifs shown in red) model by Beusch et al.[5] and MTSL rotamer clouds for sites 144 and 182; (E) same modelling with RNA removed from the model; (F) MMM simulation of the distance distributions obtained from (D) in grey and from (E) in blue; (G) CW EPR X-band spectra of doubly labelled UP1 in the presence of RNA; (H) confocal imaging of UP1 in the presence of RNA variants;

### 4. Ensemble modelling and analysis

#### a. Ensemble generation

All raw ensembles were generated in the MATLAB® software MMM[4] starting from the 20 conformers in the NMR solution state model of UP1 (pdb 2LYV,[2] residues 1-196). Residue 187 was used as the N-terminal anchor residue for the G-rich domain conformers, and the remaining C-terminal residues were removed from the starting structure (UP1 residue 1-187). The C-terminal domain was in-silico stochastically grown from the N-terminal anchor in UP1, with the appropriate amino acid sequence and assuming site-specific Ramachandran dihedral angle statistics for loop regions.[7] An independent ensemble generation run was initiated for each of the UP1 structure conformers, leading to 20 independent calculations per simulation run. The conformer generation stage of the ensemble modelling was performed with the following restraint settings: (i) no distance restraints; (ii) all 19 Gaussian fit distance distribution restrains (DDR); (iii) all possible combinations of subsets of 18 of the 19 Gaussian fit DDR, abbreviated 19-1, as described in Ritsch et al.[8]. The resulting basis sets of UP1 conformer-based hnRNP A1 conformers were then processed by ensemble refinement.

#### b. Ensemble contraction and refinement

The 20 UP1 conformer-based sub-ensembles per restraint condition (19 DDR, 19-1 DDR) were combined to the conformational basis set of full length hnRNP A1 conformers that were refined in a second stage to contract the ensemble to a smaller number of chains, while maintaining accurate representation of the experimental data. The raw ensembles comprised 2146 and 331 conformers for cases (i) and (ii), respectively. For case (iii), ensemble contraction and fitting was first performed individually for each subset of 18 restraints giving 1119 conformers from the 19 contracted ensembles. These were combined with the 60 conformers from the final ensemble obtained in case (ii) to a new raw ensemble (superensemble). At the ensemble contraction and fitting stage, we used unparametrized distance distributions obtained by Tikhonov regularization in DeerLab (using pre-release version 08b, commit ef02b20 on Jan 31 20220 at https://jeschkelab.github.io/DeerLab/). Our previous methodological study had shown only minor differences between ensembles obtained with different distance distribution fitting approaches.[8] We performed ensemble refinement with the EnsembleFit routine of MMMx (https://gjeschke.github.com/MMMx/), with the following combinations of conformational basis sets and refinement targets: (1) superensemble as basis set refined against 19 DDR+SAXS curve (DDR+SAXS ensemble), (2) superensemble as basis set refined against 19 DDR+SAXS curve + PRE data (DDR+SAXS+PRE ensemble), (3) unrestrained raw ensemble refined against 19 DDR (DDR-only ensemble) (4) unrestrained raw ensemble refined against PRE (PRE-only ensemble). We used the following terms of the target function:

*DDR (overlap deficiency):*

$$\Delta_{\mathrm{DDR}} = 1 - \left(\prod_{m=1}^{M} o_m\right)^{1/M}$$

Here, *m* indexes the *M* = 19 DDR and overlaps $o_m$ for individual DDR are defined by

$$o_m = \sum \min\{\boldsymbol{P}_{pred,m}, \boldsymbol{P}_{DDR,m}\}$$

Here, $\boldsymbol{P}_{\mathrm{pred},m}$ is the distance distribution predicted for the ensemble by rotamer modelling of the spin labels and $\boldsymbol{P}_{\mathrm{DDR},m}$ is the experimental distance distribution.

*SAXS (chi-square):*

$\chi^2$ as provided by the *crysol* routine of the ATSAS package[9]

PRE (*root mean square deviation*)[10]:

$$\rho = \sqrt{\sum_{n=1}^{N}\left[\frac{I_{para,pred,n}}{I_{dia,pred,n}} - \frac{I_{para,exp,n}}{I_{dia,exp,n}}\right]^2 / N}$$

Here, *n* indexes PRE restraints, which may correspond to the same or to different spin label sites, $I_{para,pred,n}/I_{dia,pred,n}$ is the intensity (peak volume) ratio between samples with the paramagnetic and diamagnetic label predicted by ensemble averaging and $I_{para,exp,n}/I_{dia,exp,n}$ is the corresponding experimental intensity ratio.

To balance the fitting of the different restraint subsets, we use the loss of merit *L* as a combined total fit target function:

$$L = \frac{1}{S}\sum_{s=1}^{S}\frac{m_s^{(2)}}{m_s^{(1)}} - 1$$

Here, *s* indexes the restraint subsets and the $m_s^{(i)}$ are figures of merit for the individual subsets ($\Delta_{DDR}, \chi^2, \rho$). The $m_s^{(1)}$ correspond to an ensemble fit with only subset *s* of restraints whereas the $m_s^{(2)}$ correspond to a fit with all subsets. Thus, $m_s^{(1)}$ is the lowest value that can be obtained from the raw ensemble by fitting only one type of restraints and $m_s^{(2)}$ will generally be somewhat larger, because the restraint subsets are not fully consistent with each other. Therefore, the loss of merit $L \geq 0$ quantifies inconsistency between data sets and minimization of *L* balances the relative fit qualities of the restraint subsets.[4]

The four resulting ensembles are shown in Figure S14, and the final fits to experiments are summarized in Table S4. Examples of the attained fit results for three of the ensembles are shown in Figure Figure S15 and Figure S16.

*Table S4 Figures of merit of multi-dataset model refinement*

| | DDR+SAXS[a] | DDR+SAXS+PRE[a] | PRE-only[b] | DDR-only[b] |
|---|---|---|---|---|
| PRE r.m.s.d. $\rho$ | 0.0822 | 0.0446 | 0.0581 | 0.0924 |
| SAXS $\chi^2$ | 2.016 | 2.065 | 3.108 | 3.090 |
| DDR overlap deficiency $\Delta_{DDR}$ | 0.082 | 0.092 | 0.383 | 0.179 |

[a] from superensemble as raw ensemble [b] from unrestrained raw ensemble

a. Visualization of the ensemble models

Meaningful visualization of the ensemble models of full-length hnRNP A1 with around 130 conformers can be challenging due to the disordered nature of the G-rich domain. To find a balance between accurately displaying the overall shape of the molecule and giving some intuitive understanding of the individual conformations, we used the following settings in Chimera: The anchor domains (i.e. UP1, residue 1-187) were displayed in conventional ribbon style. The G-rich domain conformers were displayed as ribbon plus solid surface at only 2 % opacity. For plots as shown in Figure S14, the transparency and white-to-red color scale of all ribbons (not the surfaces) was then individually adjusted per conformer by importing the weighting factors as a custom property with the 'Render by Attribute' tool. As opposed to main text Figure 1, the displays in Figure S14 were obtained with setting the transparency option to 'Single Layer', which better displays the individual chains in the ensemble.

b. Ensemble density analysis

The final DDR+SAXS, and DDR+SAXS+PRE ensembles were opened in Chimera software, and for each residue the command 'select *#conf_id*:*resid_reporter* za< & *r_cutoff* #0 & @CA; ' was repeated for all conformer identification numbers in the model *conf_id* (*conf_id* =0-136 for DDR+SAXS model, resp. *conf_id* =0-129 for DDR+SAXS+PRE) with the selection mode in 'append'. This selects all Cα atoms within a single conformer that are within a sphere of radius *r_cutoff* (in Å) from the residue selected by *resid_reporter*. The resulting atom identifier lists were accessed by the 'Inspect Selection' tool, and were exported for each reporter site (231, 271, 316) for spheres of *r_cutoff* = 5, 10, resp. 25 Å. Further processing was performed in MS Excel or by custom MATLAB® scripts. The average number of contacts per chain as the ratio of the total number of Cα atoms for each site and cutoff radius was obtained by dividing the total length of the list by the number of chains. This average number of contacts was further divided by the volume of the sphere of the corresponding radius, to obtain the local ensemble density in AA/$nm^3$ that is reported in the main text for the DDR+SAXS+PRE ensemble model with *r_cutoff* = 10 Å.

c. Local sequence composition analysis

The local sequence analysis was performed by using the MATLAB® 'categorical' function to make a histogram analysis of the list of amino acid residues within a sphere of chosen radius in terms of residue number and amino acid type as described above. In Figure S17 it can be seen that the choice of *r_cutoff* = 10 Å selects primarily amino acids with short (<10 AA) primary sequence distance, and contacts with remote sites occur only sporadically in individual conformers. This is what is reported in the main text for the discussion of local amino acid context bias of the reporter sites. The differences between the ensembles are weak, but note that no chain-weighting was included as an additional normalization in the histogram analysis, which might still mildly affect the histograms in the cases of large cutoff radius (> 10 Å), but is not expected to affect the local density significantly.

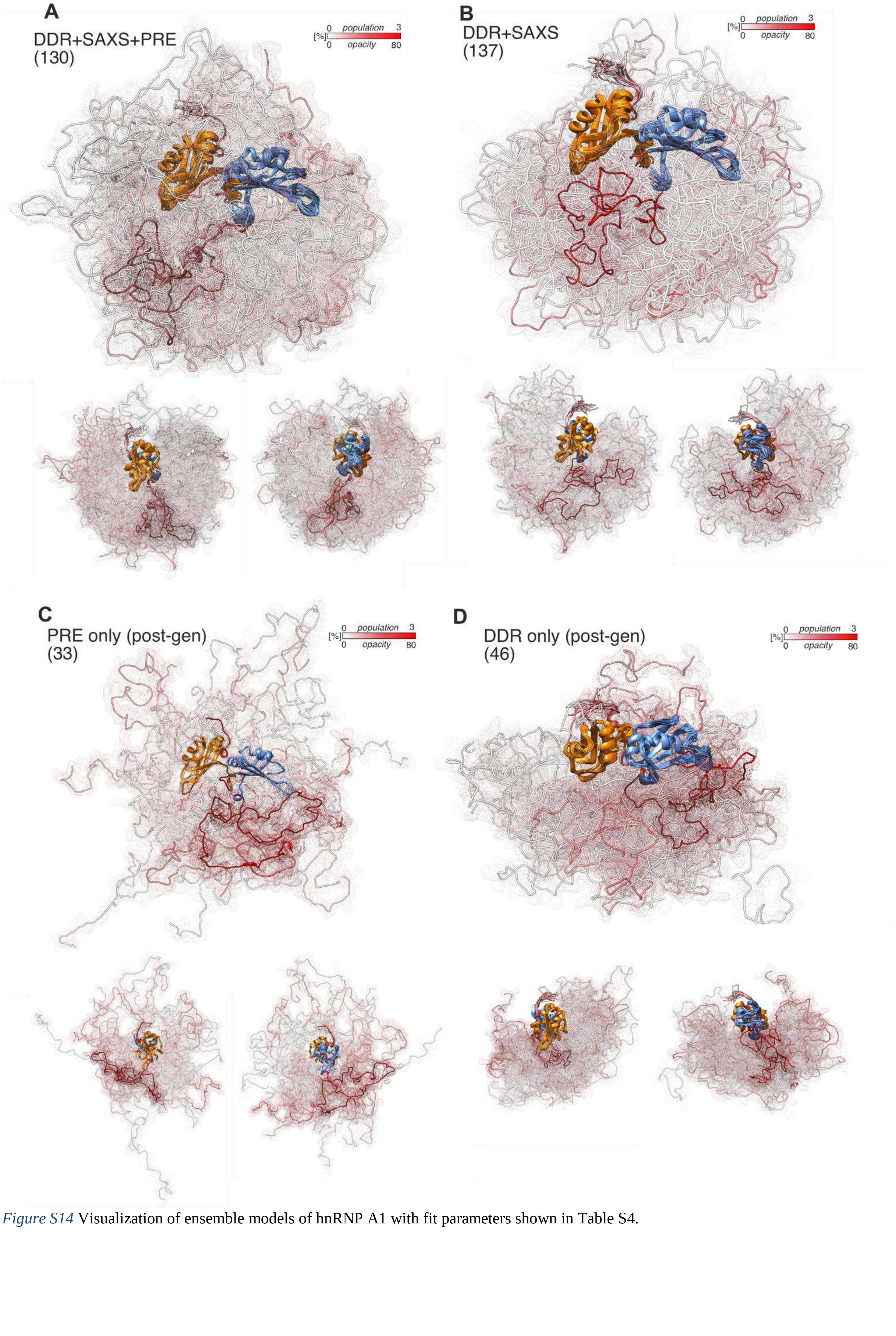


*Figure S14* Visualization of ensemble models of hnRNP A1 with fit parameters shown in Table S4.

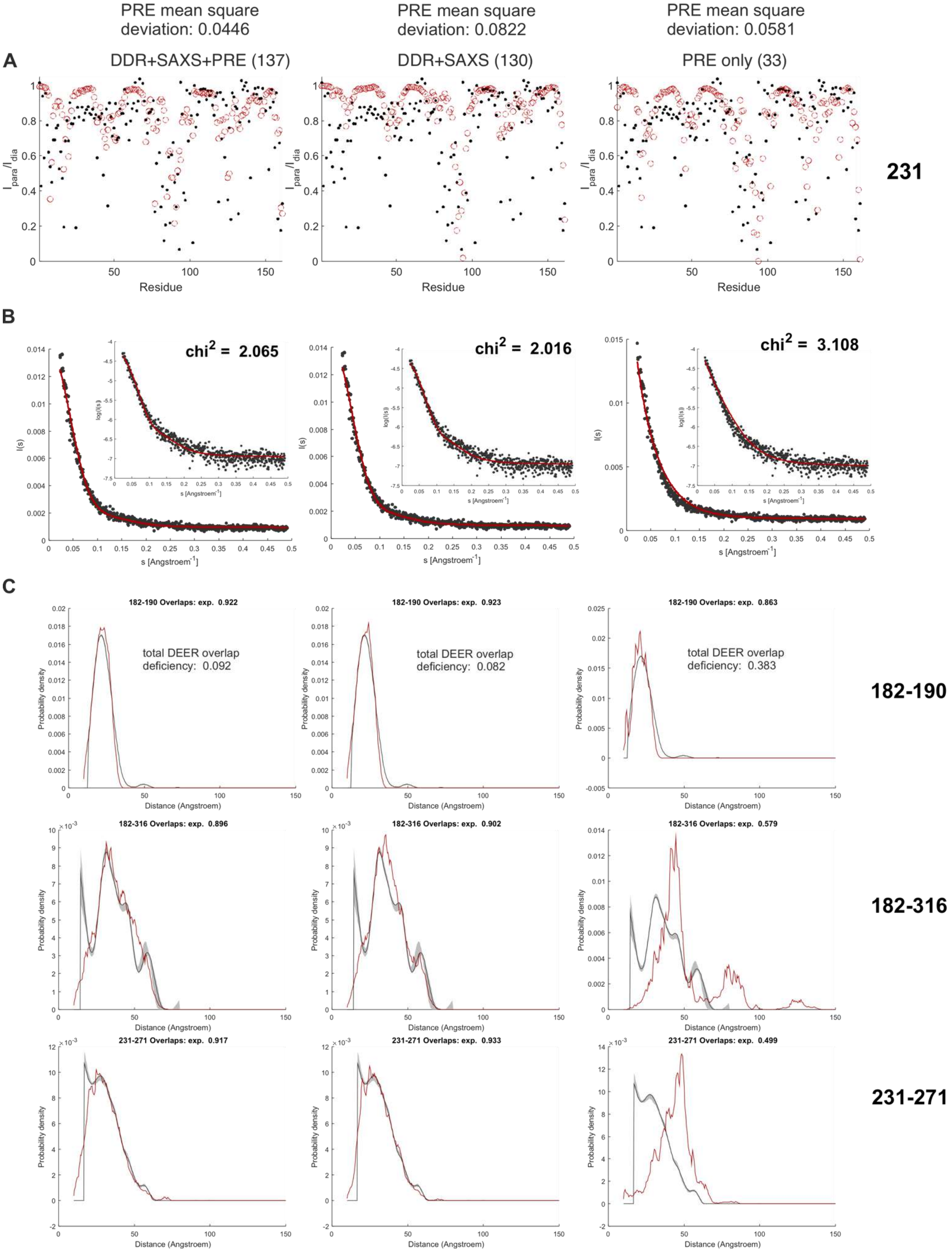


*Figure S 15* Examples of data fit targets (black) and final fulfilments (red) for (A) PRE restraints; (B) SAXS curves; (C) DEER distance restraints; when fitting only PRE data, the distance distributions are poorly reproduced.

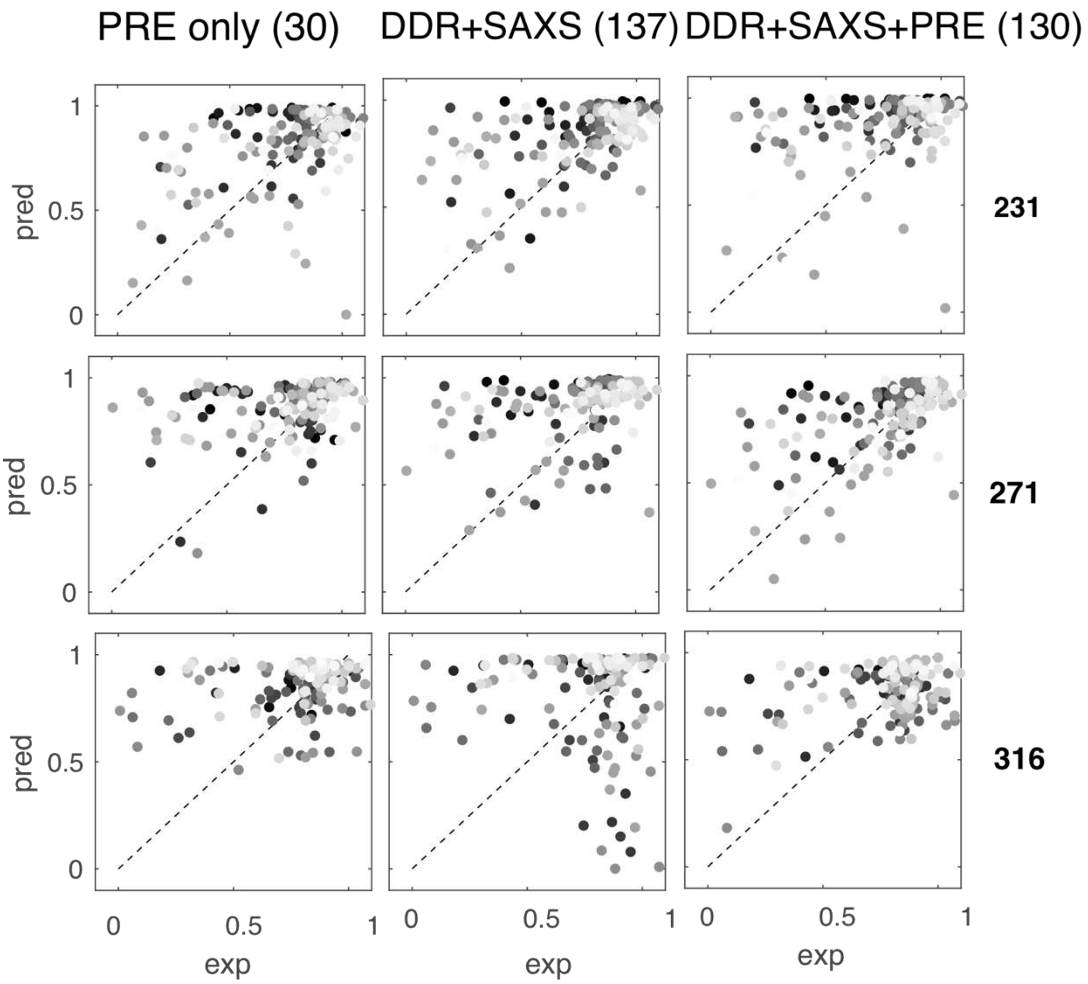

*Figure S16* PRE fulfilments for three ensemble generation schemes with full-length hnRNP A1, as summarized in Table S4.

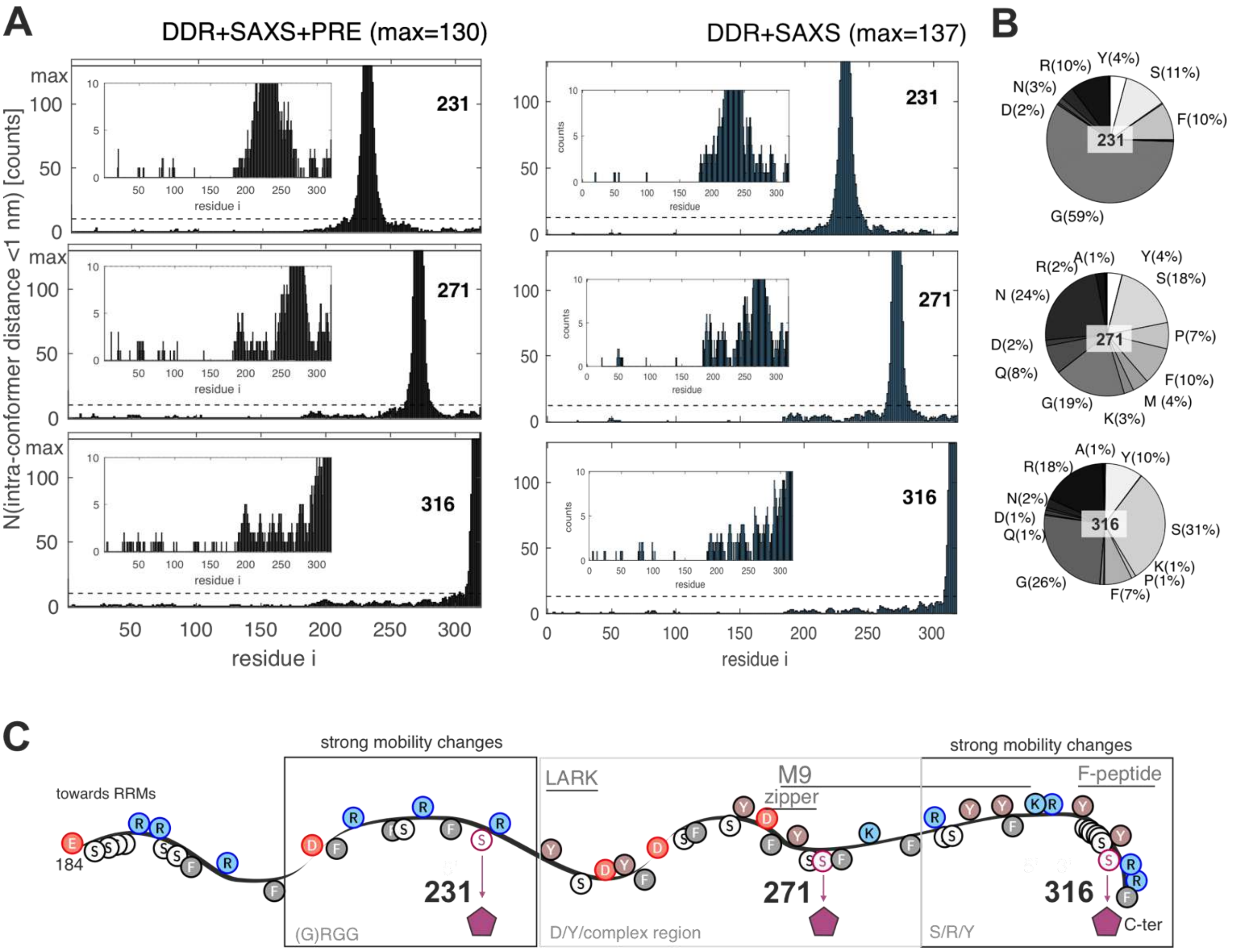


*Figure S17* Contact analysis in the ensemble model of hnRNP A1; (A) histogram of ensemble chains in which indicated residues in full length hnRNP A1 ensemble model are closer than 10 Å to the sites indicated in each panel (231, resp. 271 resp. 316); the used ensemble is indicated above (DDR+SAXS+PRE, resp. DDR+SAXS); dashed lines indicate the upper threshold in the zoomed inset plot, which shows low occupancy remote contacts (B) pie-chart of local AA abundance for reporter sites based on DDR + SAXS model; the values are very similar to those reported in main text Figure 4(D) for the DDR+SAXS+PRE ensemble (C) schematic annotation of G-rich domain of hnRNP A1 with summary of CW-EPR mobility changes upon exposure to RNA variants, and indication of some previously identified functional motifs.

### 5. Additional imaging results

#### a. Image processing

Image files were loaded into the open source Fiji software studio (https://imagej.net/ImageJ), and automated contrast enhancement was performed (default settings). Where available, z-stacks were treated the same way for all slices. For fluorescence images, the channels were split and contrast enhancement was performed only for the transmission channel. Unless stated otherwise, scale bars are 10 µm.

#### a. Additional imaging results

In Figure S18 we show additional confocal microscope imaging results, including replicates of the main imaging results with hnRNP A1 and the four ssRNAs, as well as additional imaging results obtained with mutants of hnRNP A1. We annotate some overall sample appearance observations that seem to reproducibly occur with the RNA variants, but were not further investigated in this work. In particular, the doubly mutated RNA*aa* variant seems to significantly enhance the observations of clusters of liquid droplets (LDs) that were in close contact, but not fusing. Examples are highlighted in Figure S18. Additional fluorescence results are shown in Figure S19.

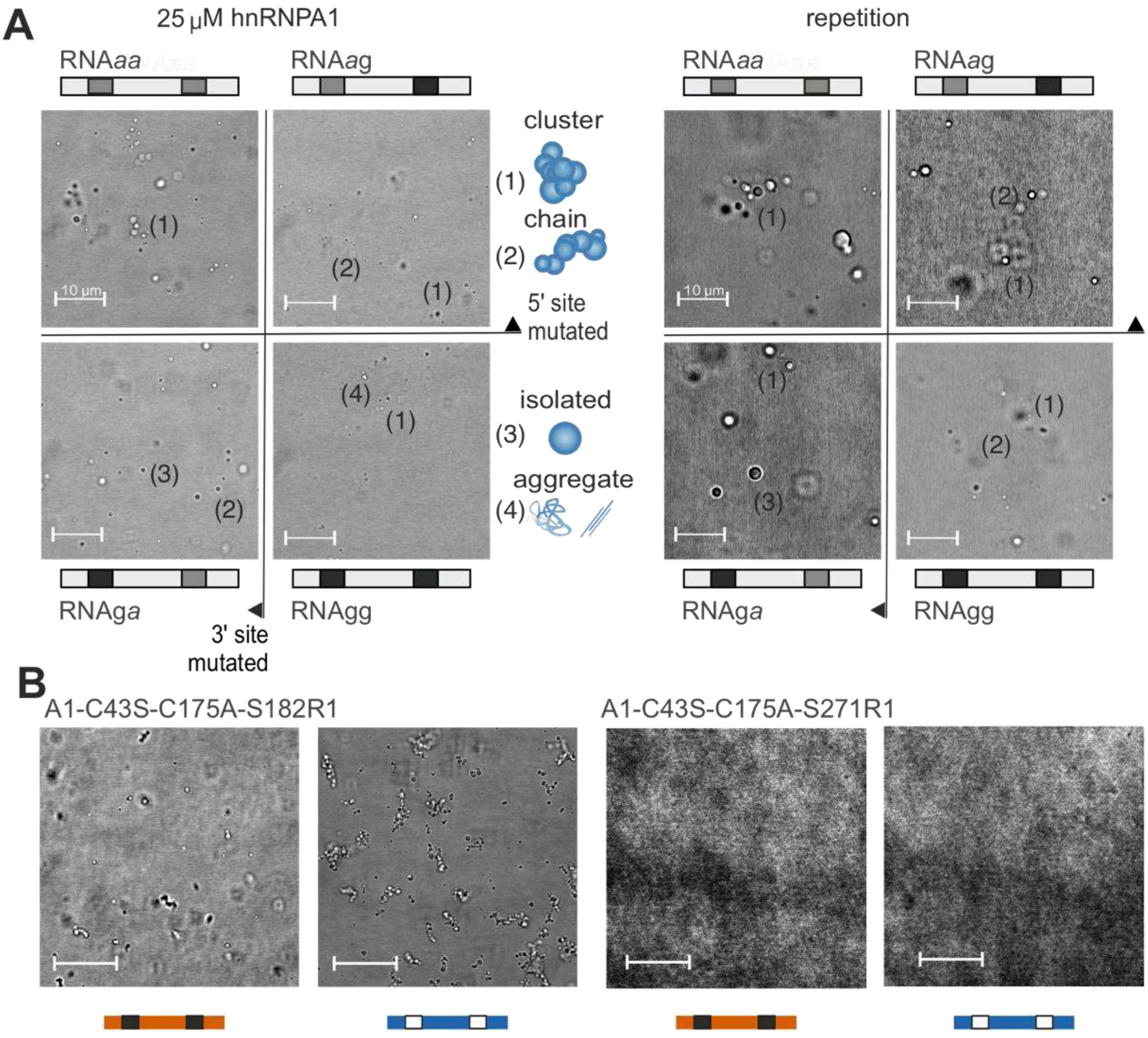


*Figure S 18* Confocal microscopy of RNA-induced LLPS of (A) hnRNP A1, resp. (B) spin-labelled mutants of hnRNP A1; clusters, chains, isolated LDs and non-spherical aggregates are indicated for some samples.

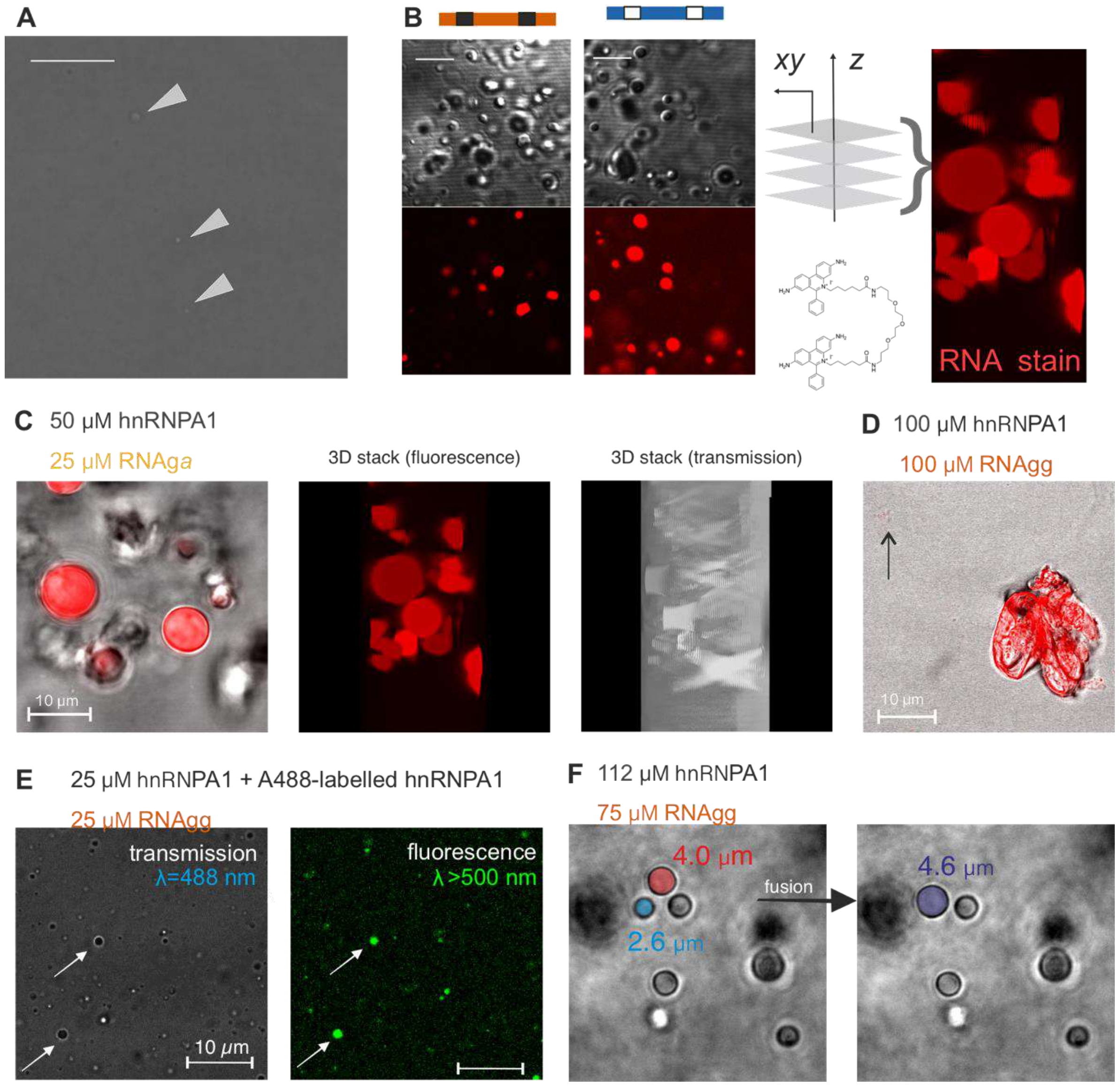


*Figure S 19* Confocal microsocpy with RNA-induced LDs of hnRNP A1 solutions; (A) transmission image of free 100 μM hnRNP A1; the very few, very small LDs that can be identified are marked; (B) schematics of deriving stacked fluorescent images of the LDs by using the RNA-specific stain 'GelRed'; (C) substochiometric addition of RNAga at high hnRNP A1 concentration led to the observation of large LDs well suited for stacked imaging; *left*: overlay of transmission (greyscale) and fluoresence channel (red); *middle*: side view of a projection from multiple z-stacks of the fluorescence channel; *right*: side view of the same projection from transmission channel; (D) overlay of transmission (greyscale) and fluorescence channel (red) of a large aggregate observed after 1:1 molar addition of RNAgg to concentrated hnRNP A1; smaller aggregates and LDs are observed simultaneously (arrow) (E) hnRNP A1 doped with unspecifically amine-labeled hnRNP A1 (NHS-Alexa-488 ); left: transmission channel, right: fluorescence channel; arrows point to same objects (F) time-course of observation of a fusion event of two large LDs that happened even in agarose-stabilised conditions; *left*: before fusion, the two LDs which are going to fuse are manually marked with colored disks, that were used to estimate of the droplet diameter (indicated); *right*: LD after fusion, with estimate of the LD diameter.

### 6. Additional PRE results

Data analysis was performed with the CARA software (Keller, RLJ. Optimizing the process of nuclear magnetic resonance spectrum analysis and computer aided resonance assignment. Doctoral thesis, ETH Zurich Thesis No. 15947, Switzerland, 2004, http://cara.nmr.ch/). The results for the all-interaction experiment with spin label at site 231 are shown in Figure S21. For better visualization of localized peak attenuation, we show projection spectra of analogous $^{15}$N-HSQC data for a repetition experiment at the same site, as well as the intermolecular-only experiment, and the all-interaction experiments at additional sites 271 and 316 in Figure S20. Integration of each assigned resonance peak in each spectrum was performed by adjusting the peak model on a well-assigned peak. We adjusted the peak list to the minor chemical shift perturbations of the resonances in the different experiments. The peak positions and the integrated peak intensities for the paramagnetic sample ($I_{para}$) and the diamagnetic sample ($I_{dia}$) were exported and analyzed with a custom MATLAB script. First, we divided each spectrum by the intensity of a reference peak (residue 117), which was selected because it is located in a region with only negligible peak volume changes between the paramagnetic and diamagnetic state. The ratio $I_{para}/I_{dia}$ was then calculated from the normalized spectral intensities and plotted as a function of the residue number. To generate the visualization on the NMR structure model, we exported the $I_{para}/I_{dia}$ table to the UCSF Chimera software.[11] The color of the surface of the UP1 solution model (pdb: 2LYV[2]) was set linearly in a blue to red colormap. The experimentally determined $I_{para}/I_{dia}$ was plotted against the predicted PRE, which is calculated as described in the modelling section above. Analogously to the main text Figure 2, for the all-interactions experiment at site 231, we mapped the intermolecular-only experiment results to the folded domains of hnRNP A1, shown in Figure S22.

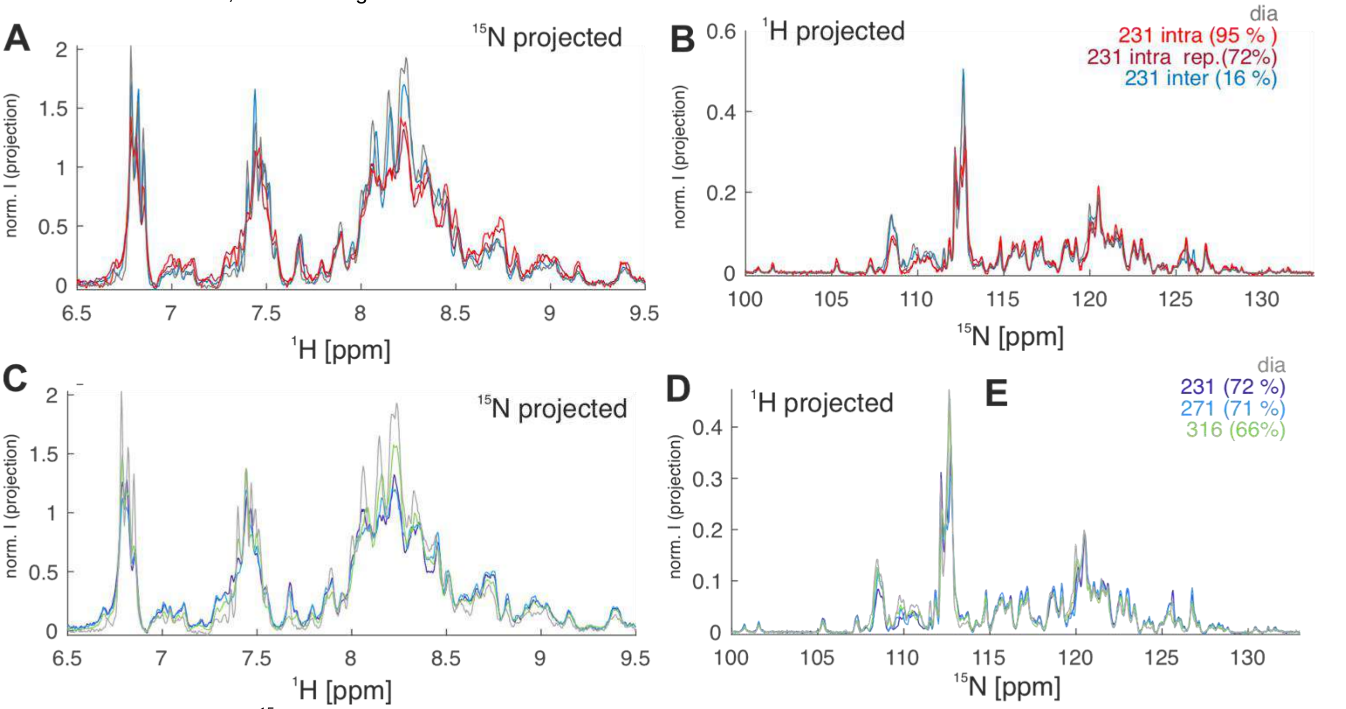


*Figure S20* Projections of $^{15}$N-HSQC spectra used for PRE analysis (A,B) repetitions of intramolecular PRE for hnRNP A1 labelled at site 231 (light and dark red), resp. intermolecular PRE (blue); (C,D) comparison for different labelling sites 231, 271, 316;

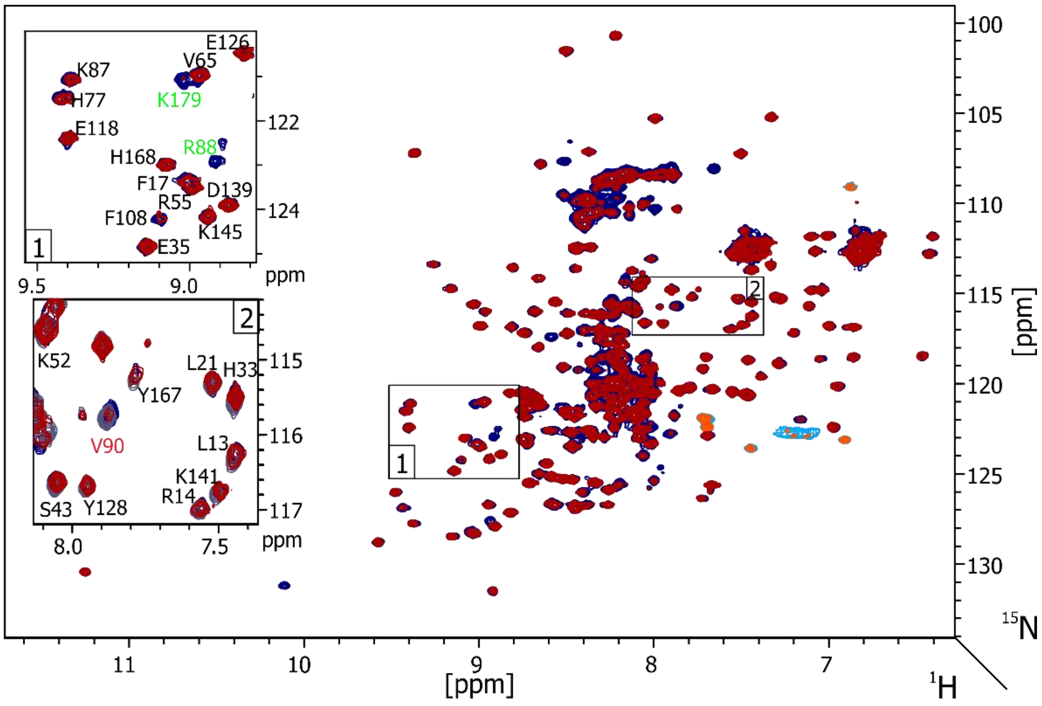


*Figure S 21* All-interactions PRE experiment with label at site 231; $^{15}$N-HSQC overlay and zoom on affected regions (insets) with labels of reference spectrum (dark blue) and paramagnetic spectrum (red).

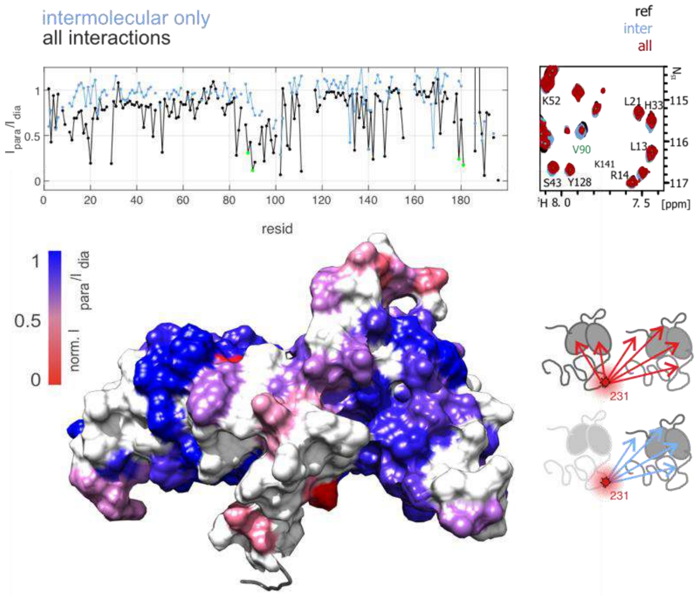


*Figure S 22* Intermolecular only, vs. all-interaction PRE experiment at site 231; (A) intensity ratio plots; (B) zoom on V90 resonance in the HSQC spectrum; (C) mapping onto UP1 solution structure; white is for AA that were not interpreted due to poor SNR in the intermolecular PRE experiment (D) schematic representation of experiment types.
